# Prodromal dysfunction of α5GABA-A receptor modulated hippocampal ripples in Alzheimer’s disease

**DOI:** 10.1101/2021.05.10.443512

**Authors:** Marcia H. Ratner, Scott S. Downing, Ouyang Guo, KathrynAnn E. Odamah, Tara M. Stewart, Vidhya Kumaresan, R. Jonathan Robitsek, Weiming Xia, David H. Farb

**Author notes:** ***One Sentence Summary***: Response of the hippocampal trisynaptic circuit to tonic disinhibition differentiates F344 from TgF344-AD rats.

## Abstract

Decades of research attempting to slow the onset of Alzheimer’s disease (AD) indicates that a better understanding of memory will be key to the discovery of effective therapeutic approaches. Here, we ask whether prodromal neural network dysfunction might occur in the hippocampal trisynaptic circuit by using α5IA as a selective negative modulator of extrasynaptic α5GABA-A receptors in TgF344-AD transgenic rats, a model for early onset AD. The results demonstrate that orally bioavailable α5IA, an established memory enhancer, increases CA1 pyramidal cell mean firing rates and peak CA1 ripple amplitude during wakeful immobility in *wild type* F344 rats resting in a familiar environment. We show that TgF344-AD rats, which express human amyloid-beta precursor protein (with the Swedish mutation) and human presenilin-1 (with a Δ exon 9 mutation), exhibit high serum Aβ42 and Aβ40 levels by 3 months of age. By 9 months of age, CA1 ripples in young adult TgF344-AD rats are nonresponsive to α5IA indicating network dysfunction prior to the onset of AD pathology and memory dysfunction. These results demonstrate, to the best of our knowledge, the first evidence for prodromal α5GABA-A receptor dysfunction in the AD hippocampal trisynaptic circuit. Moreover, as α5GABA-A receptors are located extrasynaptically and subserve the function of tonic inhibition we posit that an early stage of memory dysfunction involves the disruption of tonic inhibition in the hippocampus.

## INTRODUCTION

Forming new memories and reinforcing existing memories involves a synthesis of new sensory information with previously encoded information (*1*). Yet how neural circuitry accomplishes this function of balancing incoming sensory information represented by excitatory and inhibitory synaptic activity into memory remains a mystery. This process can be seen most clearly in the hippocampal trisynaptic circuit (HTC). In this circuit, orderly arrayed excitatory pyramidal cells (some of which function as place cells) balance excitation with inhibition in the CA3 and CA1 regions via their nearby inhibitory interneurons. The result of these concerted actions is the synchronous output of rhythmic neural activity or sharp-wave ripples (SPW-Rs) which is thought to support spatial memory in awake and resting animals (2, 3).

Hyperactivity of the HTC is associated with memory deficits such as amnestic mild cognitive impairment (aMCI) and Alzheimer’s disease (AD) in humans and animals (4–12), raising the hypothesis that increased circuit activity underlies memory dysfunction (6–8). Such a pathological increase in pyramidal cell excitability, commonly referred to as “hyperactivity”, is thought to impair memory in older humans and animals since pharmacologically reducing the hyperactivity improves cognition in amnestic mild cognitively impaired older adults (11) and aged rats (6, 12). However, paradoxically, a well-known class of memory enhancers in adult animals (13–16) (exemplified by α5IA) are thought to reduce inhibitory neurotransmission (in effect mimicking functional disinhibition) by preferentially reducing the activity of α5-containing GABA-A receptors located primarily on hippocampal pyramidal neurons (14).

To approach this apparent conundrum we asked whether reducing tonic inhibition with α5IA, which has been shown to improve memory (13–16) function, would increase CA1 place cell activity and enhance pattern separation *in vivo.* To achieve this goal, we performed *in vivo* electrophysiology experiments with a within subject design to measure real-time changes in neural network activity in the CA1 subregion as a measure of HTC output. To probe for the potential involvement of tonic inhibition as an AD-related early onset dysfunction in the hippocampal trisynaptic network we compared the dynamical responses of young adult CA1 pyramidal cells to α5IA in TgF344-AD with *wild type* F344 rats.

To the best of our knowledge, these results demonstrate, for the first time, a key modulatory role for α5GABA-A receptors in CA1 pyramidal cell mean and peak firing rates *in vivo* and ripple amplitude in adult *wild type* rats. As ripples are known to correlate with improved remembering (17) we asked whether adult TgF344-AD rats, with high plasma levels of Aβ42 and Aβ40, would also generate CA1 ripples that are sensitive to potentiation by α5IA and thus contingent upon the activity of α5GABA-A receptors. To our surprise, however, α5IA responsivity of CA1 ripples was ablated in young adult TgF344-AD rats. These findings demonstrate pharmacologically that extrasynaptic tonic inhibitory α5GABA-A receptors modulate ripple amplitude and that TgF344-AD rats exhibit prodromal disruption of this key inhibitory modulatory mechanism essential to memory circuitry function.

## RESULTS

### Reducing α5GABA-A Receptor Activity Increases CA1 Pyramidal Cell Firing Rates in Young Adult Rats

α5IA is known to cause disinhibition via a GABAergic mechanism from cell culture and hippocampal slice experiments *in vitro* (12); however, it is unknown what effect this inhibitor of α5GABA-A receptors would have on intact circuitry *in vivo* using freely moving animals (2). We hypothesized that if α5IA acted via disinhibition then it would potentiate pyramidal cell firing rates in the CA1 layer of wild type adult male Long Evans (LE) rats. To test this hypothesis, we recorded the activity of CA1 pyramidal cells using chronically implanted microelectrode arrays while rats foraged for food in a familiar environment. We used a within subject design so that each animal served as its own control. Each animal was administered vehicle followed by 3 ascending doses of α5IA **(Fig 1A).** Administration of α5IA significantly increased mean CA1 pyramidal cell mean firing rates (MFR) at the 0.3 mg/kg dose (p=0.002, two-sided signed rank test), 1 mg/kg dose (p=0.001) and 3 mg/kg dose (p<0.0001) **(Fig 1B).** Potentiation appeared to level off above 1.0mg/kg, consistent with a receptor occupancy model established previously **(Fig. 1C)** Such an increase in MFR is typically referred to as hyperactivity (6, 7) and thought to impair memory function as described above, yet α5IA is known to act as an enhancer of spatial memory in the Morris water maze (12).

**Figure 1.**
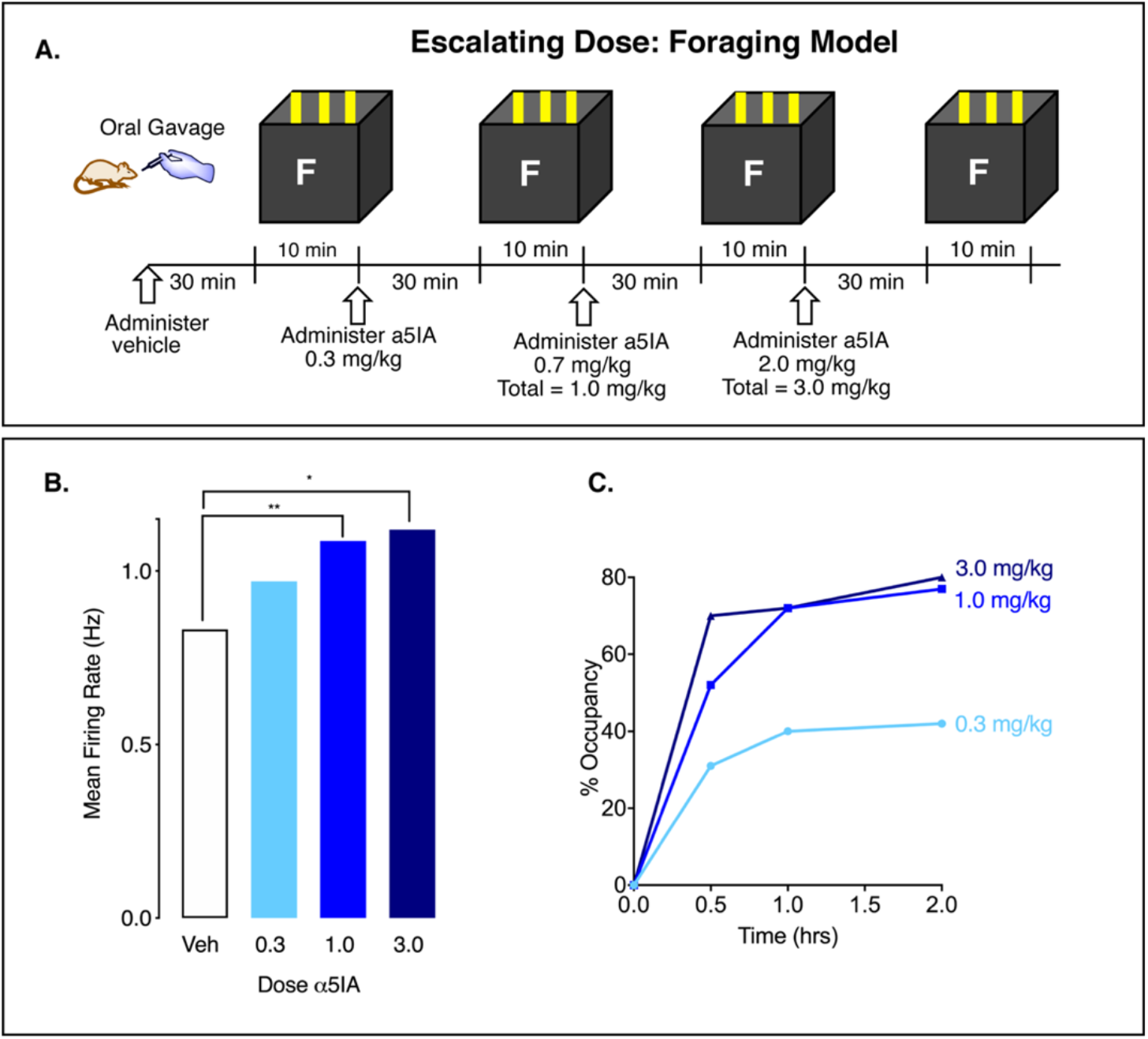
α5IA enhances LE CA1 pyramidal cell firing rates in a familiar environment. **A)** Within subjects escalating dose experiment in a foraging model. **B)** α5IA increases MFR immediately following oral administration, consistent with the dose response relationship of α5IA in panel C. **C)** Percent occupancy determined by *in vivo* dosing and *ex vivo* ligand binding levels off at 1.0mg/kg drug (data replotted from refs 11, 12). One-way repeated measures ANOVA shows dose-dependent increase in MFR (*p = 0.02; ** p = 0.003; see supplemental Table S7). Data from 4 adult male LE rats.

### α5IA-Induced Excitability of CA1 Place Cells Does Not Increase or Decrease Place Field Remapping in Adult Rats

Theories about the cellular basis for memory are rapidly evolving, with one view that subtypes of pyramidal cells such as “place cells” are differentiated from the bulk of pyramidal cells. Having established that α5IA enhances CA1 pyramidal cell mean firing rates using electrophysiology and enhances spatial memory using a behavioral test, we asked whether place cells would become hyperactive or whether they would respond differently as a function of local network activity. Lastly, we asked whether place cells would exhibit environment-dependent spatial remapping, a characteristic of the HTC believed to be a proxy for memory formation (6). Multiple single units were recorded from the CA1 pyramidal cell layer of 3 adult F344 rats (Place cell n = 152) while the animals foraged for food in familiar and novel environments (**Fig. 2A**). As anticipated, α5IA (1.0mg/kg, p.o.) increased the average mean firing rate of place cells in the familiar environment (**Fig. 2B**; planned comparisons). In the novel environment, MFR and In-field MFR increase as expected in response to novelty (**Fig. 2B**). The average peak firing rate (PFR) (**Fig. 2C**), and mean in-field firing rate (**Fig. 2D**) also increase in the familiar environment. However, no significant interaction between drug and environment was observed.

**Figure 2.**
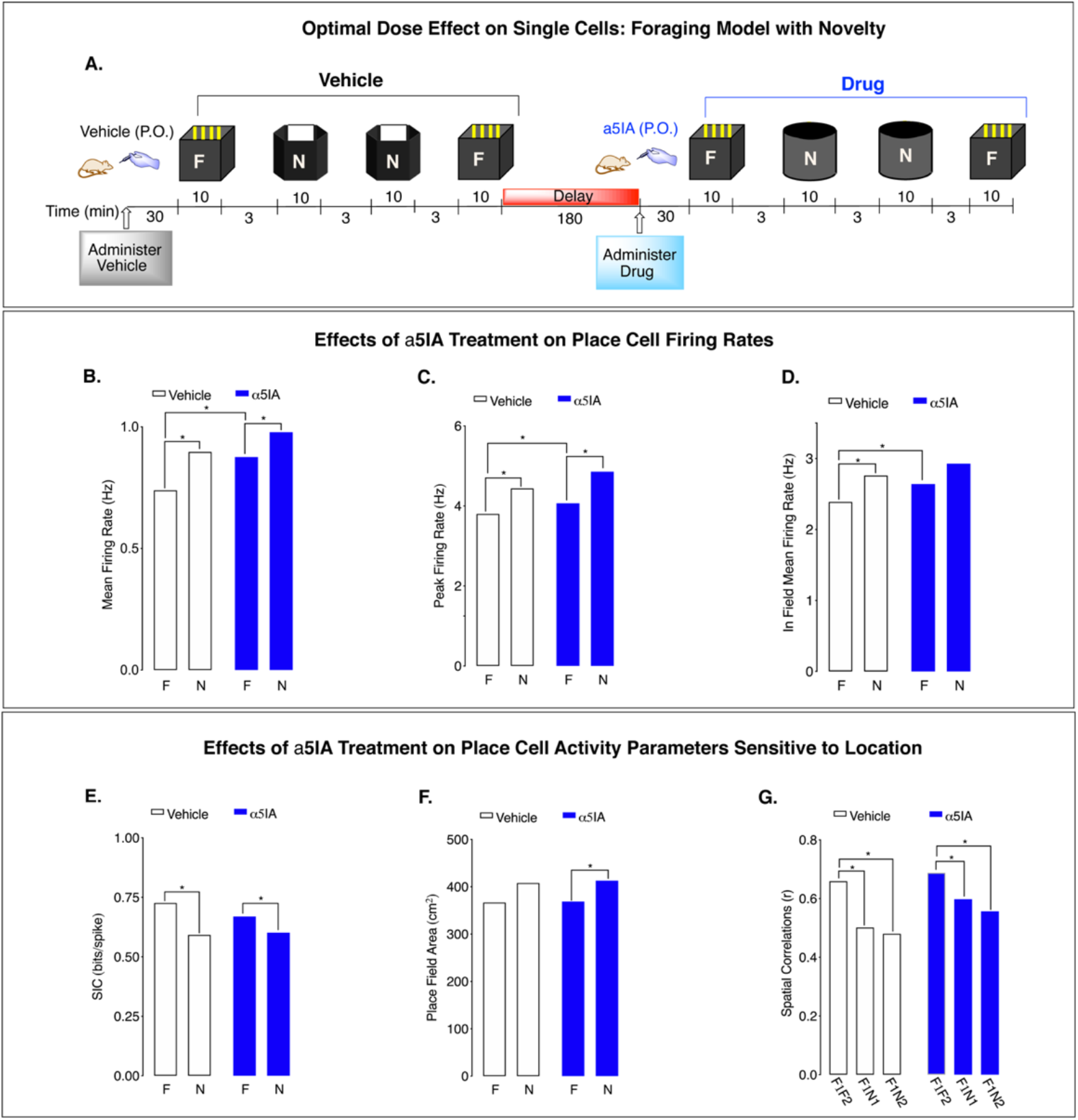
α5IA increases CA1 place cell activity in adult F344s. **A)** Environment-dependent remapping of place fields recorded across successive sessions. Administration of α5IA (1.0mg/kg, p.o.): **B)** increases MFR of CA1 place cells in a familiar environment; **C)** increases PFR in a familiar environment, **D)** increases In-field MFR in a familiar environment, **E)** has no effect on SIC in familiar or novel environments, **F)** has no effect on PFA in familiar or novel environments, and **G)** no significant effects treatment on place field spatial correlations upon exposure to environmental novelty. Significance: *p < 0.05. Data from 3 young adult F344 male rats.

Spatial information content (SIC) decreases in the novel environment, which is not affected by drug (**Fig. 2E**). Place field area (PFA) significantly increases with environmental novelty in both treatment conditions, indicating an effect of environment (**Fig. 2F**). Moreover, place field spatial correlations between the familiar and novel environments in both the vehicle and drug demonstrate that remapping occurred in both treatment conditions (**Fig. 2G**). Comparisons of spatial correlations between the two familiar environments across treatment conditions is not significant (r_veh_ = 0.77±0.12; r_drug_ = 0.76±0.13; p = 0.86), demonstrating that α5IA administration has no sig3nificant effect on the stability of well-established highly correlated place fields. The tests for interaction between drug and environment were not significant.

Despite enhancing CA1 place cell firing rates, α5IA had no effect on the spatial remapping of place cells, as quantified by the place cell spatial correlation measurement (**Fig 2G**). The SIC per action potential of place cells also was not significantly affected by α5IA (**Fig 2E**), nor was the place field size (**Fig 2F**). Surprisingly, we found no enhancement of place cell remapping by α5IA, even though we observed significant improvement on the novel location recognition test, both tests thought to evaluate spatial memory. Similar findings were observed in a group of wild type LE adult male rats **(see supplemental methods and extended data section)**.

#### Location Novelty Recognition Test Performance of Wild Type and TgF344-AD Rats

The effect of treatment with oral α5IA (1.0mg/kg) on Locational Novelty Recognition Test (LNRT) performance was assessed using a within subject design in a separate group of unimplanted LE male rats (n=9) (**Fig. 3A** **and** **B**). Vehicle LNRT was followed by a 3-hour delay in the home cage, after which the test for α5IA effects on performance was assessed. Treatment with α5IA significantly improved performance of LE rats on this task, a finding which is consistent with previous reports of this compound enhancing memory function (12, 13) **(Fig. 3C)**

**Fig. 3.**
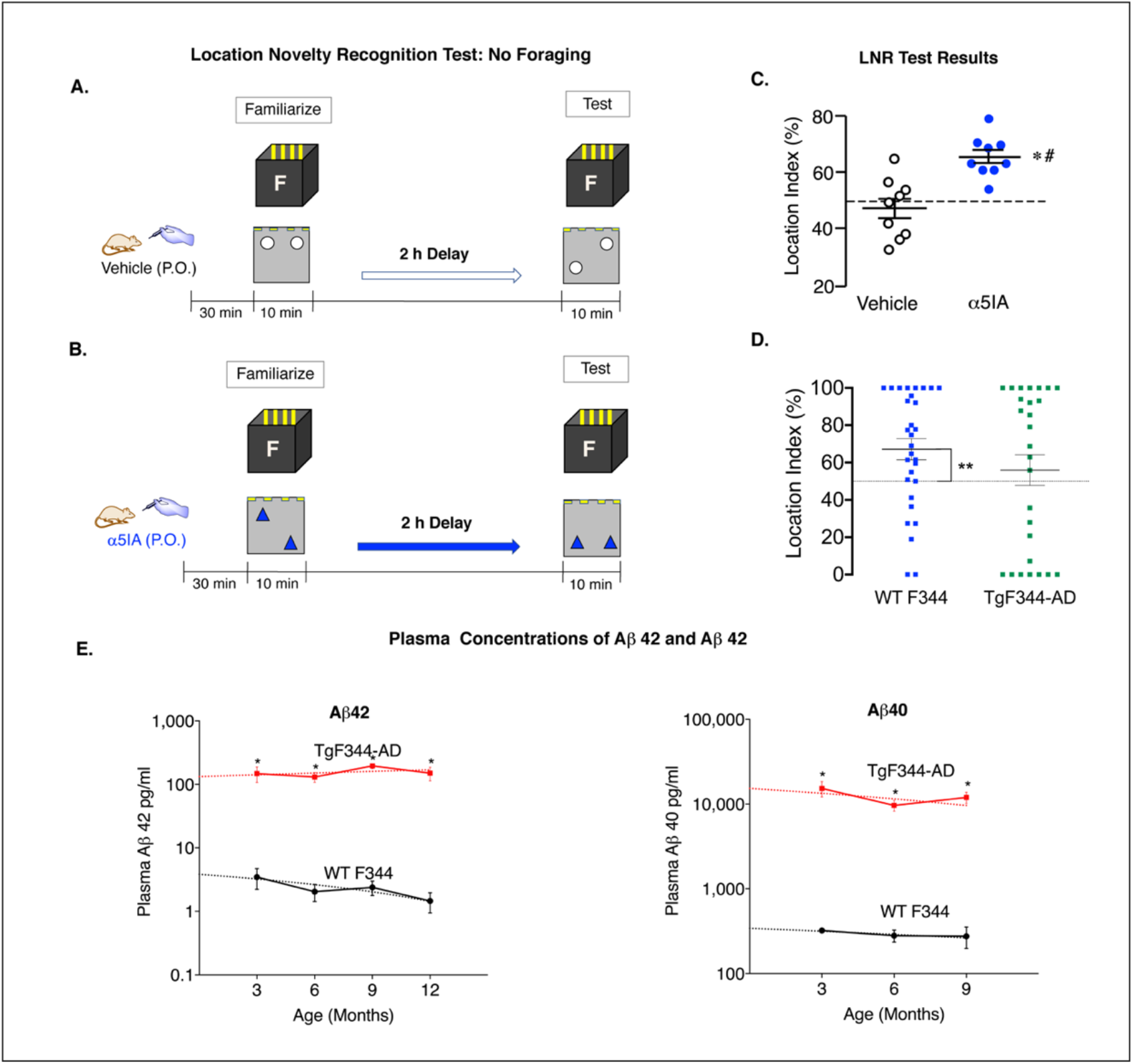
TgF344-AD rats expressing high plasma Aβ 42 levels fail to distinguish location novelty. **A & B)** Models used for assessing LNRT performance following vehicle and α5IA (1mg/kg) administration. **C)** Location index scores for individual LE rats (*n*=9) following administration of vehicle and α5IA. α5IA improved performance above chance level (dashed line; #p < 0.001) and above actual performance of at vehicle baseline (*p < 0.001). **D)** TgF344-AD rats show impaired LNRT performance. F344 rats spend significantly more time exploring objects in a novel location. TgF344-AD rats, by contrast, show no significant difference from chance in time spent exploring objects in a novel location. Significance: ** at p < 0.001. **E)** Mean plasma Aβ42 increases 4100% - 10,000% are seen at ages 3, 6, 9, and 12 mo TgF344-AD rats as compared with F344. Mean plasma Aβ40 is also increased also similarly increased at 3, 6, and 9 mo in TgF344-AD rats (green) vs F344 controls (black). Significance: * at p < 0.01.

Next the memory functions of unimplanted adult F344 (n = 30) and TgF344-AD (n = 27) rats age 9 to 18 mo, were compared in between subject design experiment. For this test, the training interval duration on the LNRT was reduced from 10 to 3 min and the delay between training and testing sessions was reduced from 2 hr to 20 min. The same previously validated objects and familiar enclosure used in the experiments assessing for α5IA effects on memory performance in LE were used in this test. The shorter training interval and delay between training and testing were selected based on previous reports indicating that F344 rats spend more time exploring the object in the novel location in this model (17). The percentage of time spent exploring the objects was calculated as a measure of memory function. The F344 rats spent significantly more time exploring the novel object location as expected. By contrast, TgF344-AD rats did not spend significantly more time exploring the object in the novel location, a finding which is consistent with impaired memory function in this rat model of AD (**Fig. 3D**). Increased plasma concentrations of Aβ42 and Aβ40 are associated with the performance deficits on the LNRT observed in TgF344-AD rats (**Fig. 3E**).

#### α5IA enhances the ripple-band power spectrum in F344 but not TgF344-AD rats

At this point we decided to focus on an aspect of HTC output as a surrogate *in vivo* biological marker of memory function. Specifically, the ripple-band frequencies (140-200 Hz) generated in CA1 and transmitted to the prefrontal cortex contain encoded information about past circuitry activity. We chose to focus on activity in the ripple band because ripples have been shown to participate in memory consolidation and are known to be coordinated by a consortium of GABAergic interneurons (2).

Based on observations from the place cell activity analyses showing no significant increases in SIC or other metrics of spatial remapping such as place field size or spatial correlations, the effects of α5IA on ripples were investigated using *in vivo* electrophysiological recordings of local field potentials (LFPs) acquired while animals were awake and immobile in a familiar environment. This behavioral model reduces theta modulation and minimizes the effects of environmental novelty and memory task demands, while at the same time promoting ripples. Specifically, we recorded LFPs from F344 and TgF344-AD male adult rats following administration of vehicle or α5IA in a paradigm that minimizes both novelty and mobilization. In F344 rats, α5IA increases ripple band power in a dose-dependent manner with the largest increase of 43% occurring at 1 mg/kg **(Fig. 4A).** By contrast, TgF344-ADs showed a decrease in ripple band power following oral administration of 1.0 mg/kg α5IA **(Fig. 4B).** Comparison of F344 and TgF344-AD for dose-response effects of α5IA revealed that the increase in ripple band power seen in F344 rats did not occur in TgF344-ADs which continue to show a pattern on serial exposure to the familiar environment after drug administration that was strikingly similar to that seen under vehicle control conditions (**Fig. 4 C-E**).

**Figure 4:**
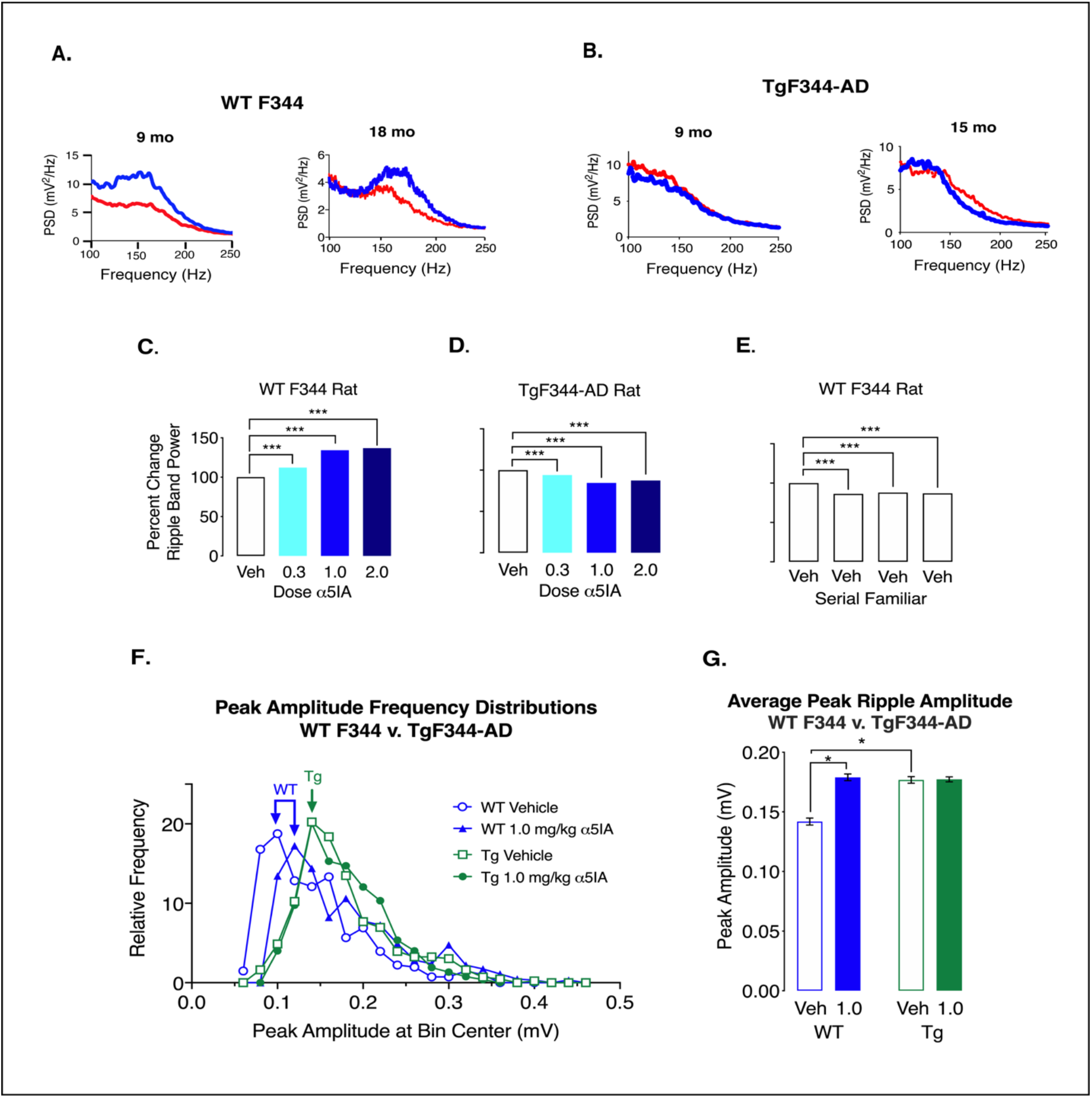
Administration of α5IA does not increase ripple band power or peak ripple amplitude in TgF344-AD Rats. **A)** Visual comparison of the spectrums **(top panel)** from representative well-positioned electrodes in two F344s age 9 & 18 mo reveals a remarkable overt increase in ripple band power following administration of α5IA (1.0 mg/kg). **B)** Visual inspection of the power spectrums from well-positioned tetrodes adult TgF344-AD rats age 9 to 15 months demonstrates a decrease in ripple band power following administration of α5IA. **C)** Histogram showing percent increase in ripple band power in F344 rats following administration of escalating doses of α5IA. **D)** Histogram showing percent decrease in ripple band power in TgF344-AD rats following administration of escalating doses of α5IA. **E)** Percent decrease in ripple band power in adult F344 rats following administration of vehicle only during serial exposures to a familiar environment. **F)** Peak ripple amplitude frequency distributions in F344 before and after oral administration of 1.0 mg/kg α5IA (n = 3) show a shift to the right that is not seen in TgF344-AD rats (n = 3). **G)** Peak ripple amplitude significantly increases in F344 following 1.0 mg/kg α5IA. In addition, peak ripple amplitude is significantly greater under vehicle conditions in TgF344-AD rats as compared with F344s. Significance: * at p < 0.05, ** p < 0.01, and *** p < 0.001.

Visual inspection of the frequency distributions for ripple event peak amplitudes reveals skewed distributions in both F344 and TgF344-AD subjects **(Fig. 4F).** α5IA shifts the frequency distribution to the right in F344 but not TgF344-AD animals. This is consistent with the observation that α5IA increases the mean peak amplitude of ripples in F344 but not TgF344-AD rats (**Fig. 4G**). Interestingly, TgF344-AD genotype increases the average peak ripple amplitude at baseline to a level that matches the effect of α5IA on F344s. It is tempting to propose that this increase in CA1 ripple amplitude may be a compensatory prodromal response to early, potentially memory impairing, effects of circulating beta-amyloid.

## DISCUSSION

The current observations in TgF344-AD (as compared with F344) rats reveal a significant prodromal increase in CA1 peak ripple amplitude that is nonresponsive to α5IA, consistent with a homeostatic prodromal response to early, potentially memory impairing, effects of circulating beta-amyloid. The present findings are associated with significantly elevated plasma concentrations of Aβ42 and Aβ40 in TgF344-AD rats, implicating bioaccumulation of Aβ as the likely active principle in neural circuitry dysfunction.

This hypothesis of prodromal disruption of HTC network function in AD is consistent with the finding that ripple amplitude predicts which information is remembered (18) and that disruption of SPW-Rs during rest impairs spatial learning (19). When non-human primates perform memory-guided visual search tasks the SPW-Rs associated with trials that are “remembered” have larger amplitudes than do the SPW-Rs associated with trials that are “forgotten” (18). By contrast, the duration of SPW-Rs does not change based on whether a trial is remembered or forgotten. These results establish a relationship between SPW-R amplitude and which trials are remembered. Chronic circuitry level hyperactivity due to the loss of functional GABAergic interneurons has been proposed to underlie memory dysfunction associated with age-related mild cognitive impairment (20). Yet acute administration of levetiracetam plus valproic acid rapidly reverses the memory impairment in aged rats (*12*) and chronic hyperactivity as measured by decreased mean firing rate, increased spatial information content, and place field sharpening to be indistinguishable from young adults (6). These observations suggest that the HTC may very well be morphologically intact but physiologically dysfunctional.

In conclusion, these findings indicate that plasma Aβ burden increases peak ripple amplitude and a decrease or loss of tonic extrasynaptic α5GABA-A receptor mediated control over ripple dynamics in the CA1 subregion. To the best of our knowledge this may represent the first investigation of α5GABA-A receptor function and dysfunction in AD and of ripple modulatory function as a possible prodromal marker for AD onset *in vivo*.

## Supporting information

Supplemental Methods and Experimental Data

## ACKNOWLEGEMENTS

We dedicate this paper in memory of our collaborator Howard B. Eichenbaum, PhD, who was instrumental in helping us establish *in vivo* electrophysiology as a platform for advancing an understanding of the pharmacology of memory. His scientific contributions to this research sadly ended prior to completion of the research. We also would like to thank Sam McKenzie, PhD, and John Bladon, PhD, for their many thoughtful and helpful comments and suggestions over the past two years. DHF thanks Michaela Gallagher (John’s Hopkins) for her generous contributions toward helping us establish this line of investigation.

## Non-author Contributions

The authors thank Dr. L. Adrienne Cupples for her assistance with statistical analysis and Dr. Yunlong Bai of the Harbin Medical University in China for his assistance with our validation of the offline ripple detection code used in these experiments.

## Author contributions

TMS, HBE (sadly not listed as an author as deceased), RJR, and DHF conceived of and designed the place cell and behavioral experiments; MHR and DHF conceived of and designed the sharp wave ripple experiments; MHR, OG, KEO, and TMS performed experiments; OG implanted animals; TMS and VK performed histology; SSD wrote analysis code in MATLAB and performed computer data analysis; MHR and SSD analyzed data; and MHR, SSD and DHF wrote the manuscript.

## Data and Materials Availability

All data needed to evaluate the conclusions presented in this paper are present in the paper and/or the supplementary materials section. The datasets and the custom MATLAB scripts used in this study are available upon request.

