## Supplemental Methods and Experimental Data for "Prodromal dysfunction of α5GABA-A receptor modulated hippocampal ripples in Alzheimer’s disease"

### Title: Prodromal dysfunction of $\alpha$5GABA-A receptor modulated hippocampal ripples in Alzheimer's disease

**Supplementary Methods and Extended Data**

**This section includes:**

Methods
Supplementary data
Figs. S1 to S18
Table S1 to S15
Citations to References 21–32

#### METHODS

##### *Subjects*

*The in vivo* electrophysiological experiments reported herein were performed on n = 20 adult male rats. Escalating dose experiments looking at pyramidal cell firing rates during exploration of a familiar environment were performed in four wild type male LE retired breeders ages 8-12 mo. Studies of CA1 place cell remapping activity in response to environmental novelty were performed in three wild type LE male retired breeders ages 8-12 mo and three wild type F344 males ages 8-12 mo. Investigations of dose-dependent effects of $\alpha$5IA on ripple activity when animals were immobile in a familiar environment were performed in 1 LE male rat age 15 mo; 4 F344 rats ages 9-18 mo; and, 3 adult male TgF344-AD rats ages 9-16 mo. Two 11 mo old male F344 rats and one 11 mo male TgF344-AD rat were used for the vehicle control experiments.

The novel location recognition behavioral experiments were performed in a separate group (*n* = 9) of male LE retired breeders age 8-12 mo that had not undergone surgical implantation of microelectrode arrays.

All rats were individually housed in a climate-controlled vivarium maintained on a regular 12hr/12hr light/dark cycle in the Laboratory Animal Science Center at the Boston University School of Medicine. Rats had *ad libitum* access to water but were mildly food deprived to 85% of their free-feeding weight during training and testing. Rodent housing and research were both conducted in strict accordance with the NIH Guide for the Care and Use of Laboratory Animals. Boston University is accredited by the Association for Assessment and Accreditation of Laboratory Animal Care. The Boston University Institutional Animal Care and Use Committee approved all procedures described in this study.

##### *Drugs*

The GABA-A $\alpha$5-selective positive allosteric modulator $\alpha$5IA (3-(5-methylisoxazol-3-yl)-6-[(1-methyl-1,2,3-triazol-4-yl) methyloxy]-1,2,4-triazolo[3,4-a] phthalazine) (Sigma Aldrich Inc, USA and Tecoland, Irvine, California, USA) was chosen for these studies due to its unique pharmacodynamic profile. Oral administration of $\alpha$5IA to rats at doses of 0.3, 1.0 and 3.0 mg/kg have all been shown to enhance hippocampus-dependent memory in the absence of locomotor, anxiogenic, or epileptogenic side effects (*12*). Memory enhancing effects of α5IA occur when the drug is administered 30 minutes before trial commencement at which time receptor occupancy for the 0.3, 1.0 and 3.0 mg/kg doses were determined to be 25%, 55%, and 68% respectively (*11*). The variance about the occupancy values in the 0.3 mg/kg dose was over three times that of the 1.0 mg/kg and 3.0 mg/kg doses (*11*). The intermediate dose of 1.0 mg/kg was chosen for these *in vivo* electrophysiology studies because the occupancy percentage and variance at this dose are more favorable than those associated with a lower dose and, because the higher dose did not significantly improve performance above that shown with the 1.0 mg/kg dose on the Morris water maze (*11-13*) This dose was also predicted have fewer off-target effects than the higher dose.

The vehicle (0.5% 400 centipoise methylcellulose) and drug were both administered via oral gavage 30 min before initiation of testing. To facilitate the oral administration process, the rats were mildly sedated by placing them in an induction chamber filled with 5% isoflurane. The drug was always administered after vehicle; counterbalancing the order of vehicle and drug administration was not possible in this model due to the duration of the experimental protocol, the plasma half-life (0.9 hr) of the drug and, our desire to record the effects of α5IA, and vehicle from the same cells in the animals on the same day to minimize electrode drift.

##### *Microelectrode Arrays*

Custom microelectrode arrays constructed with a cast polymer core (Smoot-Cast 300; Smooth-On Inc., Easton, PA) and fitted with 24 independent micromanipulators permitted positioning of individual nichrome tetrodes (Sandvik, Palm Coast, Florida, USA) within the brain region of interest. Each micromanipulator consisted of a 0.5 in, 23 gauge stainless steel tube affixed with cyanoacrylate adhesive to a Delrin bridge. A flexible fused silica capillary tube (Polymicro Technologies™), was inserted into this 23-gauge guide tube and secured into position with cyanoacrylate adhesive to provide structural support for the tetrodes. A 0-90 x 0.5 in brass machine screw (J.I. Morris, Southbridge, Massachusetts, USA) was attached to a Delrin bridge and used to facilitate vertical movements of the microdrive. Twenty-four stainless steel 30-gauge guide tubes were inserted in the polymer core to provide additional support for the silica capillary tubes. These tubes where bundled together at the bottom of the microarray by insertion into a 1.0 cm piece of 14-gauge stainless tubing. The entire bundle of guide tubes was then sealed with dental acrylic to create a de facto mounting point for subsequent surgical implantation of the completed microarray. Each tetrode was comprised of four nichrome wires (California Fine Wire, Grover Beach, CA, USA); the tips of which were gold plated to decrease impedance to 200 kΩ at 1 kHz.

The F344 and TgF344-AD rats used in the vehicle control experiments were all implanted with Buzsaki 32 type silicon probe microelectrode arrays (NeuroNexus, Inc. Ann Arbor, Michigan, USA) affixed to a custom micromanipulator.

##### *Surgical Procedures*

Rats were anesthetized with isoflurane (3.5% for induction; 1.5% - 2% thereafter) in 100% oxygen delivered via a calibrated vaporizer (Vaporizer Sales & Services Inc., Rockmart, Georgia, USA). Buprenorphine (0.05 mg/kg, s.c.) was administered 30 min before surgery to provide intraoperative analgesia. Glycopyrrolate (0.02 mg/kg, subcutaneous; s.c.) was also administered to reduce salivary and bronchial secretions and prevent vagal bradycardia. The heads of the animals were shaved and prepped for surgery with betadine in triplicate. After placing the rats in a stereotaxic instrument (David Kopf Instruments, Tujunga, CA), a midline sagittal incision was made across the top of the head to expose the cranium. A small craniotomy (~2 mm in diameter) was created immediately above the right dorsal hippocampus at 3.6 mm AP and 2.6 mm ML from bregma. The dura mater was excised and the electrode array lowered into position above the surface of the neocortex. Ten additional holes (< 0.5 mm) were drilled into the skull for the placement of an equal number of stainless-steel screws; two of these screws served as electrical grounds as well as providing additional anchoring points for securing the microelectrode array to the skull with dental acrylic (Patterson Dental Supply, St. Paul, Minnesota, USA). At the end of surgery, electrodes were advanced ~750 μm into the cortex. The craniotomy was sealed using Kwik-Sil silicone sealer (World Precision Instruments, Shanghai, China) for tetrodes or a warm melted mixture of parrafin wax and mineral oil for slilicon probes. Post-operative analgesia was maintained for 72 hr with buprenorphine (0.05 mg/kg, s.c.). Lactated ringers (6ml every 12 hr), soft food, and Hydrogel (ClearH2O, Westbrook, Maine, USA) were also administered during the first 72 hr after surgery to ensure adequate caloric intake and hydration. Topical triple antibiotic and Cephalexin (60 mg/kg, p.o.) were administered for seven days post-operatively to reduce the risk for infection at the implant site.

***Data Acquisition and Spike Sorting***

Recordings of place cell and LFPs were acquired from custom nichrome tetrode microarrays or Buzsaki 32 channel type silicon probes (NeuroNexus, Inc. Ann Arbor, Michigan, USA) using a 96 channel Plexon Multichannel Acquisition Processors (Plexon Inc., Dallas, TX) and an Intan RHD system respectively. Recordings were referenced to a common skull screw for LFPs, or an indifferent tetrode (for single unit activity), and hardware filtered for LFPs (0.77 – 400 Hz) and spikes (154 Hz – 8.8 kHz). LFP signals were amplified 1000X and digitized at 1 kHz while spike signals were amplified 2000 – 8000X and digitized at 40 kHz. The location of the rat within each environment for recordings of place cell activity was tracked in real-time with an overhead camera imaging two LEDs connected to the headstage (HST/32V-G20-2LED) at a rate of 30 frames/sec (Cineplex, Plexon Inc., Dallas, Texas, USA). The appearance of theta activity during ambulation and ripple events in the LFPs when animals were immobile was used to functionally confirm electrode location within the CA1 hippocampal subregion. If pyramidal cell activity was not identified on an electrode during daily screening, it was advanced and allowed to settle for at least 1 hour before further assessment for single unit activity.

Shareware (KlustaKwik) developed by Ken Harris (*21*) was used to sort action potentials into clusters based on three principal components using an expectation-maximization algorithm. Only well-defined clusters with clear refractory periods having an interspike interval greater than 1ms were included in the final analysis. Clusters from pyramidal cells were differentiated from interneurons based on firing rates, waveform widths (≥ 300 μs for pyramidal cells; < 300 μs for interneurons) and the presence or absence of a burst of activity in the 3-10 ms range on inspection of the auto-correlograms (*21*).

***Escalating Cumulative Dose Paradigm for Analysis of CA1 Hippocampal Pyramidal/Place Cell Activity***

Rats were allowed to explore a black plywood square (60 cm x 60 cm) enclosure with vertical yellow stripes on one wall for at least 10 days to establish this as a "familiar environment". The environment was centered on a table with solid black impervious floor which surrounded by blackout curtains to mask background cues. The floor of the environment was cleaned with 30% ethanol between session to remove olfactory cues. Crushed fruit-flavored cereal crumbs were randomly distributed over the floor of the recording chambers to encourage continuous ambulation and exploration of the entire environment. The dosing schedule used in this model was selected based on previously reported benzodiazepine receptor occupancy over time following oral administration of α5IA (11-13). To achieve total cumulative *in vivo* systemic doses of 0.3 1.0 and 3.0 mg/kg, the drug was administered via the oral gavage 30 minutes before the beginning of each recording session at concentrations of 0.0, 0.3, 0.7 mg/kg and 2.0 mg/kg. This cumulative dosing schedule combined with a within-subject repeated-measures design facilitated recording the dose-response curve for CA1 pyramidal cell (n = 60) neural activity from the same cells on a single day during four serial 10-minute recording sessions acquired while animals foraged for food in the familiar environment. Between sessions rats were placed in holding container (30 cm x 30 cm x 45 cm) which was gently rotated for ~3 min to disorient the animals immediately before placing them into the next environment. Rats had the opportunity to drink water between environmental exposures. *(6, 22, 23).* Only data from animals that showed good coverage of all four environments during was included in final analysis (**Fig. S1**).

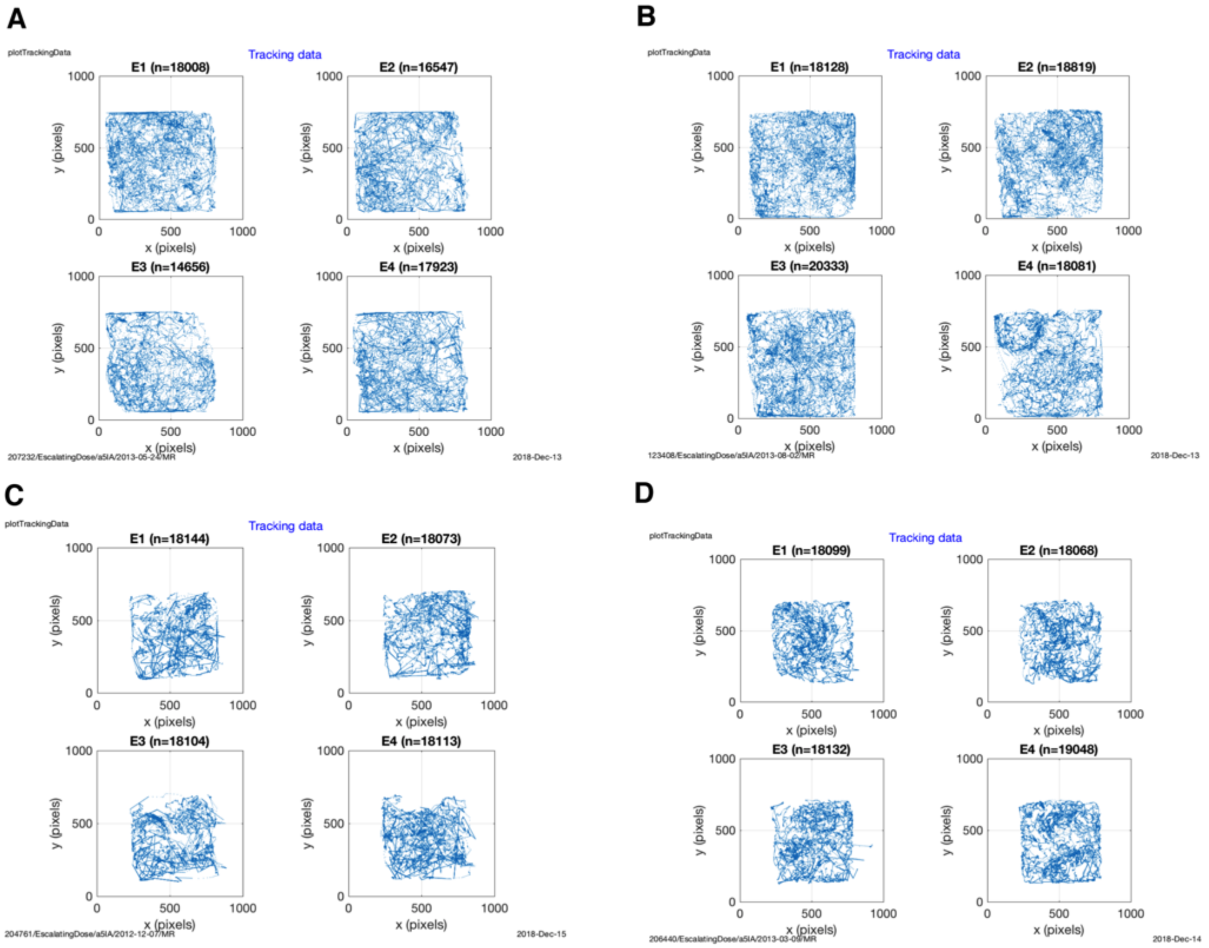


**Fig S1: Trajectories of Long Evans Adult Male Rats During Escalating Dose Model.**

***Environment-Dependent Place Cell Remapping Paradigm***

During the training phase of the remapping experiments, rats were again allowed to explore a black plywood square (60 cm x 60 cm) enclosure with vertical yellow stripes on one wall for at least 10 days to establish this as a "familiar environment". During the test phase, place field remapping was assessed as the animals explored a "novel environment" (hexagon or cylinder with the same pattern of vertical stripes) in addition to the now familiar environment in the following order: Familiar–Novel–Novel–Familiar **(Fig**

**S2).** All other procedures were identical to those described above for the escalating dose experiments performed in the familiar environment.

##### *Place Cell Firing Analysis*

Place cell activity was recorded from three LE male rats while the animals explored the Familiar–Novel–Novel–Familiar environments. Pyramidal cells in the CA1 hippocampal subregion that fired at least 60 action potentials during each of the 10-minute recording sessions (minimum rate 0.1 Hz) were analyzed for place fields using a series of custom MATLAB (Mathworks, Natick, MA) scripts developed in our laboratory. Occupancy-normalized spatial firing rate maps from animals showing good coverage of the environments based on trajectory data **(Fig. S2 and S3)** were estimated using the total number of action potentials (spikes) occurring in a given bin divided by the total amount of time spent in the bin (3.5 cm x 3.5 cm), with distinct firing rate maps calculated for each exposure to the familiar and novel environments. The smoothed value for each bin was calculated as the mean for each bin, and all bins within 5 cm, where each bin was weighted by its distance from the central bin using a two-dimensional Gaussian kernel (*24*). Bins included in the analysis must have been visited at least once during the session. Place fields were defined as six adjacent bins with averaged binned rates above 0.5 Hz and averaged binned firing rate > 2x mean overall firing rate (all bins). Data was only included for spikes that occurred when running speed was > 2.0 cm/sec. For each environment, the pixel-to-pixel correlations, and the spatial selectivity were calculated only if the overall mean firing rate was > 0.1 Hz.

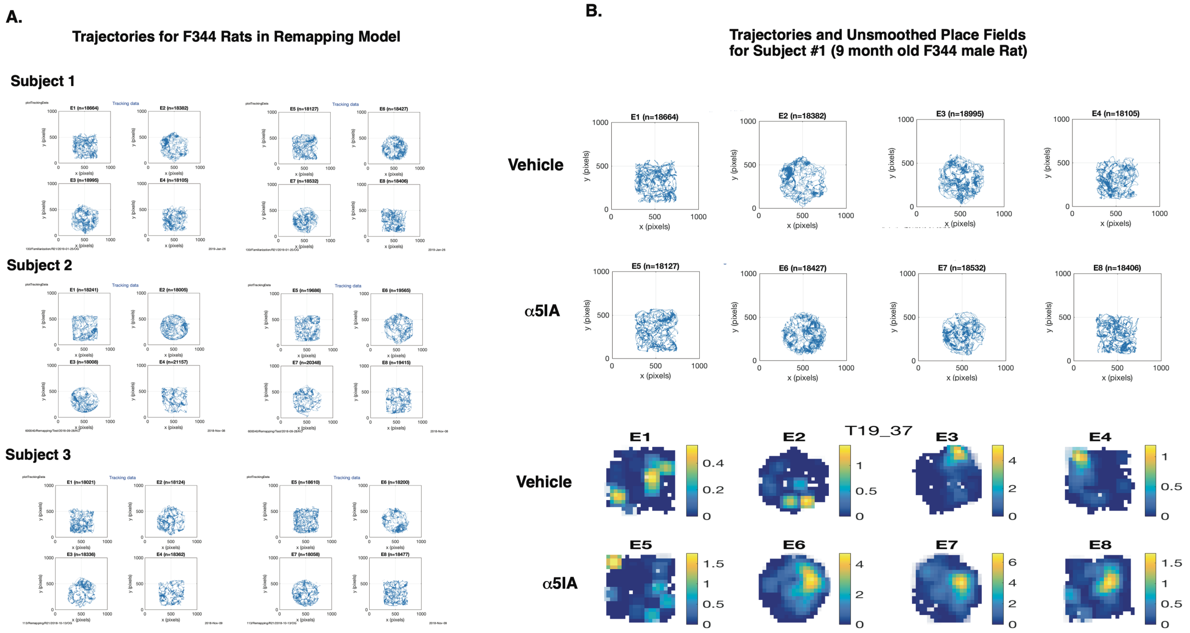


**Fig S2. A)** Representative trajectories of F344 male rat during place cell place field remapping experiment. **B)** Representative path trajectories (top) and unsmoothed place fields (bottom) from a remapping experiment in a F344 rat (subject 1) showing increased firing rates (rate remapping) due to novelty under vehicle control conditions and following oral administration of α5IA (1.0mg/kg). Figure also shows that the overall mean firing rate is also higher following α5IA administration in both environments as compared with vehicle.

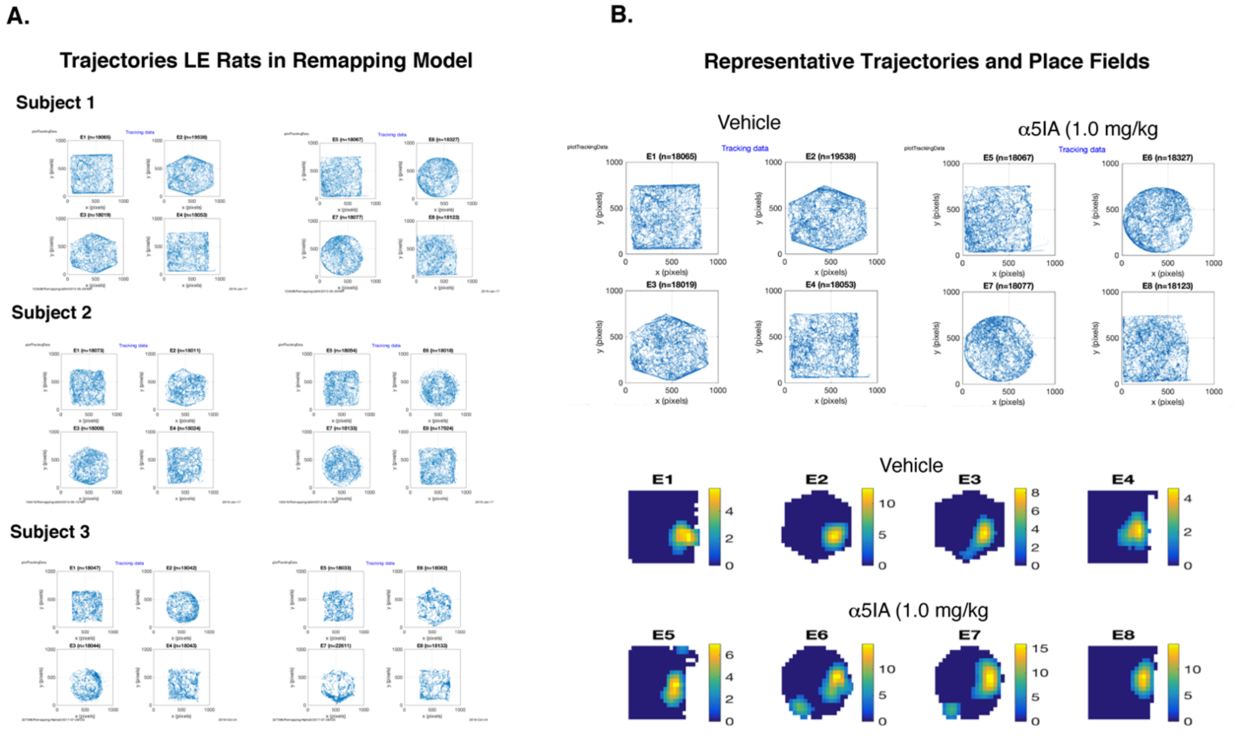


**Fig. S2: A)** Trajectories from LE Rats during Place Cell Remapping Model. **B)** Representative trajectories (top) and unsmoothed place fields (bottom) from Subject 1 showing characteristic increases in firing rates due to novelty and oral α5IA (1.0mg/kg) administration.

***Place Field Analysis***

To evaluate the effects of environment and treatment with α5IA on place cell characteristics we applied measures to assess for discharge frequency (mean and peak firing rates). Selectivity and stability of place cell firing, across environments under the two treatment conditions was also determined by calculating the 1) place field area, 2) spatial information content, 3) and spatial correlations, for each place cell.

1) PLACE FIELD AREA was computed for each environment as the sum of all bins that met the place field criteria. Prior to this computation, the firing rate maps for the novel environments were transformed to a square shape to facilitate comparison with the familiar environment (*25*).

2) SPATIAL INFORMATION CONTENT provides a measure of the amount of information each spike conveys about the animal's position in the environment (in units of bits/spike) (*26*).

$$Information\,per\,spike = \sum_{i=1}^{n} P_i\left(\frac{\lambda_i}{\lambda}\right)\log_2\left(\frac{\lambda_i}{\lambda}\right)$$

where $i$ = 1,…, $n$ is the bin number, $P_i$ is the probability of occupancy at bin $i$, $\lambda_i$ is the mean firing rate for bin $i$, and $\lambda$ is the overall mean firing rate of the cell.

3) SPATIAL CORRELATIONS were evaluated by computing the bin-to-bin Pearson correlation between environments (F1 v. F2; N1 v. N2; F1 v. N1). Confidence intervals of the correlations were determined after applying the Fisher z transform which were then transformed back and reported as the as means ± SEM (*27*). Previous studies have shown that place fields in the CA1 subregion are not only environmentally specific, but that once established, these place fields are very stable (28). Based on these previous findings, only place cells with well-correlated place fields (r > 0.5) in the familiar environment demonstrating that these place fields were stable under both treatment conditions were used to asses for the effects of environmental novelty on place field remapping based on changes in spatial correlations.

$$z = \frac{1}{2}\, ln\left(\frac{1-r}{1+r}\right)$$

***Cell by Cell Analysis for Rate Remapping***

Individual CA1 place cells recorded from LE male rats were defined as showing rate remapping if the average in-field mean firing rate change in the two novel environments was 50% or more above or below the average in field mean rate in the two familiar environments.

***Escalating Cumulative Dose Paradigm for Analysis of Ripples***

A cumulative escalating dosing schedule was used to assess for within subject effects of $\alpha$5IA on ripples in the CA1 hippocampal pyramidal subregion of adult male LE, F344 and TgF344-AD rats to minimize the influence of environmental novelty on ripple events, serial 10-minute recordings of LFPs were acquired while animals were awake and immobile in the familiar environment Food was withheld to discourage ambulation. **(Fig S3 and Table S1**). Localization of electrodes was based on visual examination LFP oscillations for distinct oscillations (e.g., high-frequency ripples events and low-frequency sharp waves) and observing synchronous activity across recording sites **(Fig. S4).** Because our initial experiments in the LE animal did not reveal a significant increase in full session ripple band power as compared as with vehicle following the 3.0 mg/kg dose, the decision was made to reduce the maximum dose to 2.0 mg/kg in all subsequent experiments in the F344 and TgF344-AD rats. The cumulative dosing concentrations were adjusted to 0.0, 0.3, 0.7 and 1.0 mg/kg to achieve total cumulative *in vivo* systemic doses of 0.3, 1.0 and 2.0 mg/kg. Recordings were made for 10 min while rats were immobile resting quietly in one location of a familiar environment. Because electrode depth can influence power in ripple band, the electrodes were not moved during these recordings **(see Fig S4B and C).** With exception of food being withheld to discourage ambulation all other procedures were identical to those described above for the *escalating*-dose firing rate analysis experiments which were also performed in the same familiar environment.

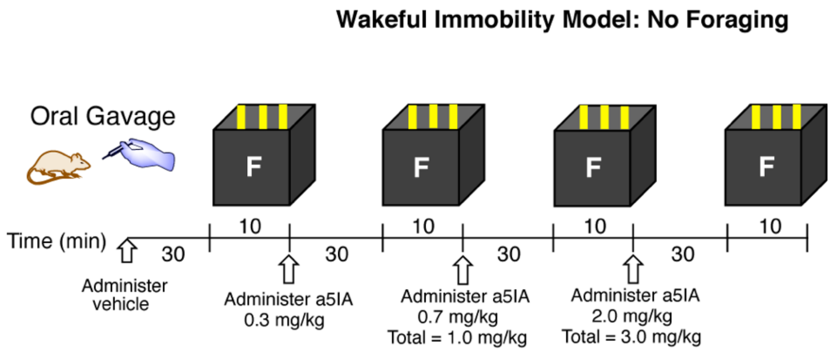


**Fig. S3. Detection of CA1 ripples. A)** Schematic diagram of LFP recordings for analysis of vehicle and drug effects of ripples. Data was acquired from immobile awake animals placed in a familiar environment. Food was withheld to discourage foraging behavior during the 10 min recording sessions.

**Table S1. Ages of Strains of Rats Used in Escalating Dose Experiments for Probe Drug effects on Ripples**

| **Subject/Rat #** | **Strain** | **Age at testing** |
|---|---|---|
| Subject 1/#327396 | LE | 15 |
| Subject 2/#130 | F344 | 9 |
| Subject 3/#600040 | F344 | 16 |
| Subject 4/#142 | F344 | 18 |
| Subject 5/#113 | F344 | 12 |
| Subject 6/#115 | TgF344-AD | 12 |
| Subject 7/#128 | TgF344-AD | 9 |
| Subject 8/#822 | TgF344-AD | 11 |
| Subject 9/#811 | F344 | 11 |
| Subject 10/#814 | F344 | 11 |

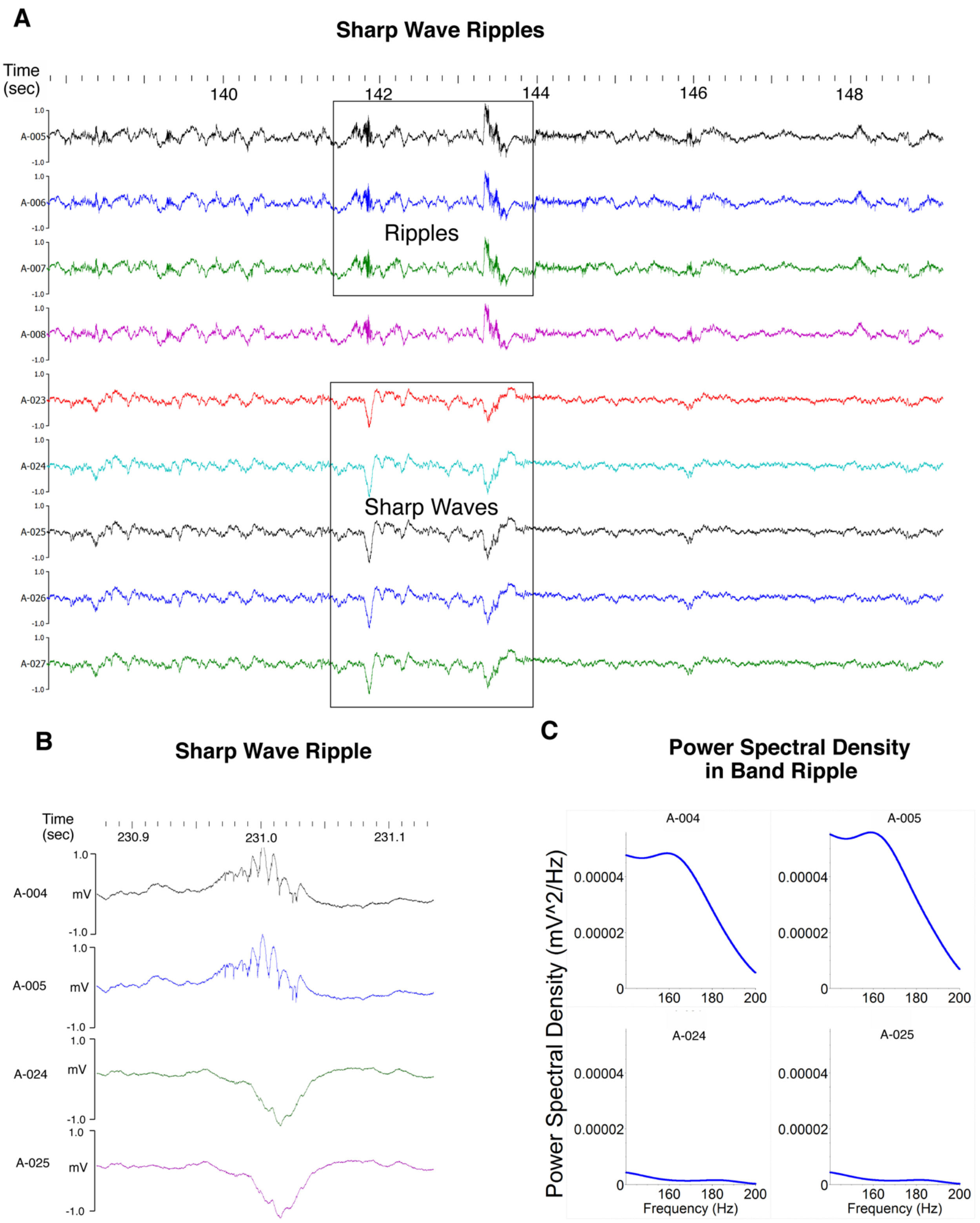


**Fig S4. Sharp waves ripples. A)** Distinct depth dependent characteristic oscillations (e.g., high-frequency ripples events and low-frequency sharp waves) and synchronicity of activity across recording sites is seen on visual inspection. **B and C).** LFP and power spectrum for sharp wave ripple recorded with two electrodes showing greater power in ripple band on the two electrodes that detect ripple and lower power on the two detecting the associated sharp wave.

***Local Field Potential Signal Processing***

Prior to statistical analysis, LFPs were frequency filtered to capture activity in the 100-250 Hz frequency band and plotted visual presentation of overt differences in ripple band (140 to 200 Hz) power at each dose. The effects of α5IA on peak ripple amplitude, peak ripple frequency and ripple duration were quantified using the LFP data from the electrode best positioned in the CA1 pyramidal cell layer at vehicle baseline. Electrodes were not moved between test sessions. Data was filtered for periods of wakeful immobility based on the delta (1.5-4 Hz) to theta (4-10 Hz) power ratio (29). **(Fig. S5 and Table S2).** To identify ripples, the LFP signal voltage was bandpass filtered (140-200 Hz), squared and normalized. A custom MATLAB script validated against manually curated sharp wave ripple events was used to detect ripples. To ensure that selected events were genuine ripples, only events that deviated from the background voltage by more than 5 standard deviations (SD) at their center were defined as ripple centers (30). Ripple boundaries defined by deviations greater than 2 SD. To further ensure that ripple detected were not false positive events, we determined the single unit firing times relative to the onset time of all ripples **(Fig S6).** Ripples with inter-ripple spacing less than 30 ms were merged, while those events shorter than 20 ms or longer than 100 ms excluded. The dose dependent change in raw power for each FFT frequency was defined as the dependent variable of intertest (n) and compared with itself across all 4 doses tested. All PSD data are reported in $mV^2/Hz$ and analyzed using a within subjected repeated measures ANOVA.

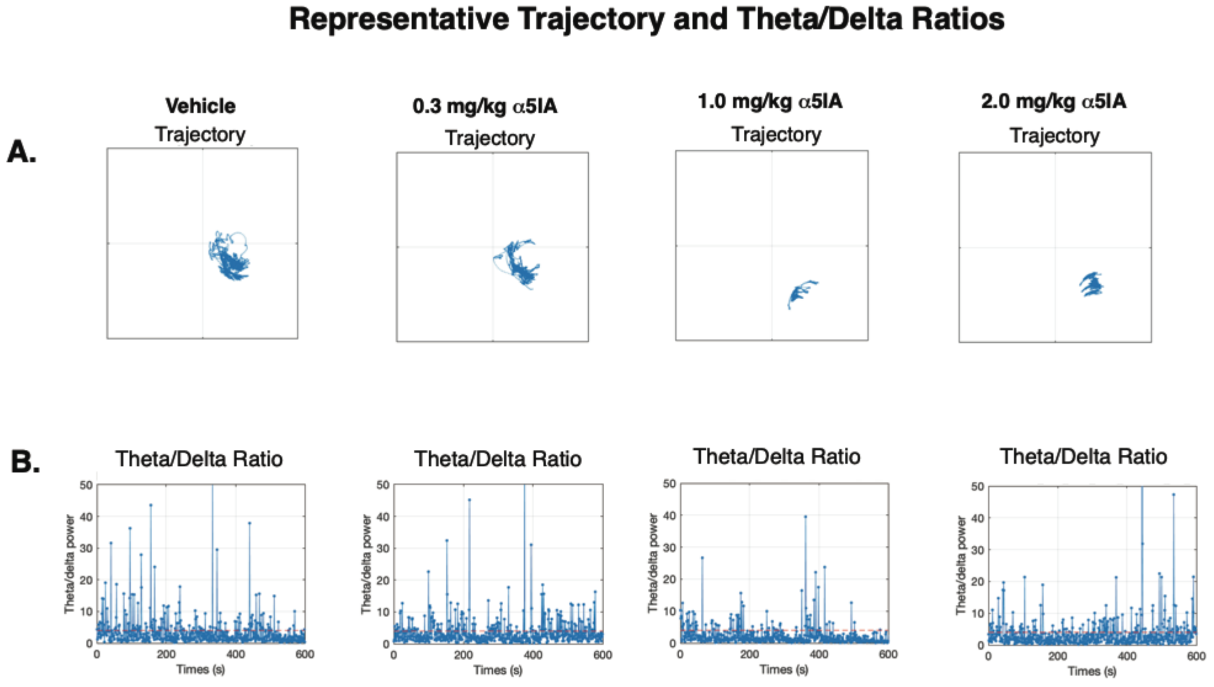


**Fig. S5: Representative trajectories and theta/delta power ratio.** All data shown are from Rat #2 (9 mo old F344 male). Panel A shows trajectories and panel shows corresponding theta/delta power ratios during serial exposures to the familiar environment in the escalating dose model.

**Table S2: Percentage of Time Spent Immobile During Test Sessions**

| Rat/Drug/Dose/Model | Percent Time Immobile |
|---|---|
| LEM_Subject-1_Veh_FFFF | 49.003 |
| LEM_Subject-1_α5IA_0.3_FFFF | 50.332 |
| LEM_Subject-1_α5IA_1.0_FFFF | 50.831 |
| LEM_Subject-1_α5IA_3.0_FFFF | 43.76 |
| F344 _Subject-2_Veh_FFFF | 77.434 |
| F344 _Subject-2_α5IA_0.3_FFFF | 74.799 |
| F344 _Subject-2_α5IA_1.0_FFFF | 85.578 |
| F344 _Subject-2_α5IA _2.0_ FFFF | 77.449 |
| F344_Subject-3_Veh_FFFF | 84.242 |
| F344_Subject-3_ α5IA_0.3_FFFF | 83.233 |
| F344_Subject-3_ α5IA_1.0_FFFF | 94.591 |
| F344_Subject-3_α5IA_2.0_FFFF | 89.583 |
| F344 _Subject-4_Veh_FFFF | 85.548 |
| F344 _Subject-4_α5IA_0.3_FFFF | 84.653 |
| F344 _Subject-4_α5IA_1.0_FFFF | 87.725 |
| F344 _Subject-4_α5IA _2.0_ FFFF | 83.845 |

#### Relationship Between Ripples and Multi-Unit Activity

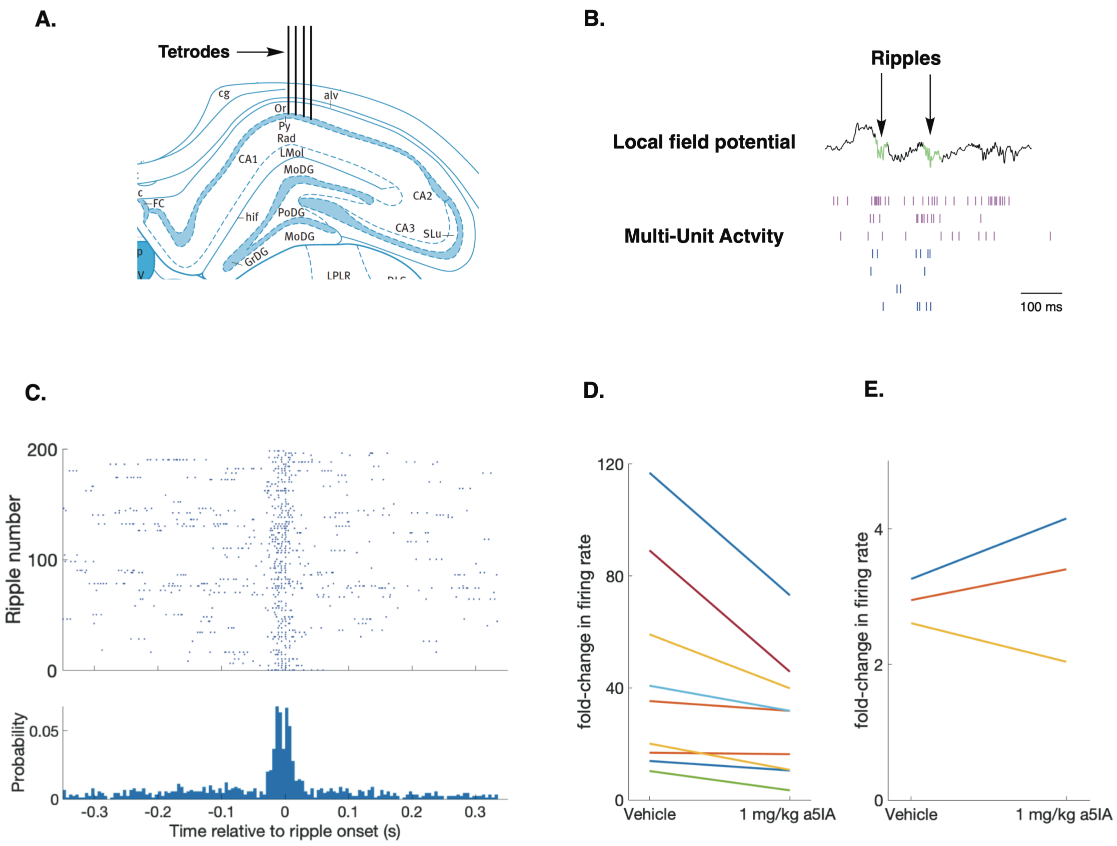

**Fig S6: Single unit firing times relative to the onset time of ripples.** **A)** Schematic drawing of representative recording sites in the hippocampal CA1 subregion. **B)** Representative multiunit activity during ripples. **C)** Ripple-triggered raster plot (top) and firing probability histogram (lower) for a representative CA1 pyramidical cell. Fold-change in ripple-associated firing rate with and without $\alpha$5IA for (**D**) pyramid cells and (**E**) interneurons recorded with electrode best located within the pyramidal cell layer.

***Histology***

At the end of the study, rats were deeply anesthetized with pentobarbital (100 mg/kg; i.p.) and the recording electrode locations marked by passing anodal current (30 $\mu$A, 5 sec) through each wire of the tetrodes. The animals were then perfused with ice cold phosphate buffered saline (7.4 pH) followed by either neutral buffered 10% formalin or fresh 4% paraformaldehyde. Brains were extracted and post-fixed at 4C, cryoprotected in 15 and 30% sucrose. All brains were inspected for and found to be devoid of lesions and tumors. 50$\mu$M sections were obtained and Nissl stained prior to microscopic inspection and imaging. **(Fig. S7)**

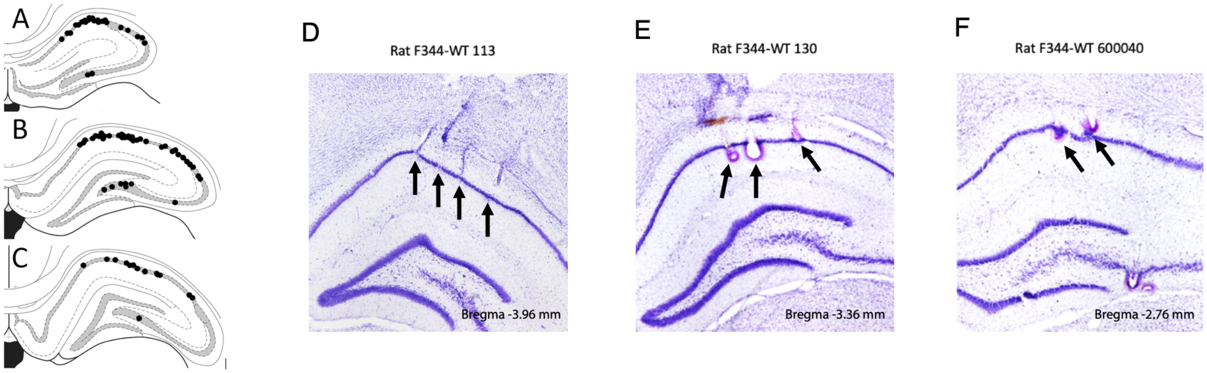


**Fig. S7. Histological verification of recording locations.** Illustration of coronal sections that are in the anterior-posterior plane at -2.5 mm (***A***), -3.4 mm (***B***), and -3.8 mm (***C***) from Bregma for rats used in place field remapping studies. Black dots indicate tetrode locations. Majority of tetrodes were located in CA1 and those that were located elsewhere were excluded from analyses. Photographic images of coronal sections **(D, E, F)** showing examples of tetrode locations in CA1 subregion for three F344 male adult rats used for ripple analysis.

***Quantification of Plasma A$\beta$***

Blood samples (200-250 ml) were taken from the tail veins of F344 (and TgF344-AD rats. Plasma samples (100 ml) were prepared using 10% potassium K-EDTA mixed with whole blood at final 0.1% K-EDTA dilution. Blood sample were centrifuged at 1500g to extract supernatant plasma. Prepared plasma samples were stored at -80°C for subsequent biochemical analysis with ELISA. A Meso Scale Discovery (MSD) platform was used to test the samples. Assays for Aβ 40 and 42 were analyzed using V-PLEX Aβ peptide Panel 1, following the manufacturer's instructions (MSD, Gaithersburg, Maryland, USA). An analysis of the variance with Kruskal-Wallis test for plasma concentrations of A$\beta$42 and A$\beta$40 in F344 and TgF344-AD revealed no significant differences due to sex. Data from males and females was therefore merged prior to analysis for effects of genotype on A$\beta$42 and A$\beta$40 (**Table S3**).

**Table S3: Aβ42 and Aβ40 Plasma Concentrations in Males and Females.**

| **Aβ42** | **Multiple comparisons test Adjusted P Value** |
|---|---|
| TgF344-AD 3M v. 3F | >0.9999 |
| TgF344-AD 6M v. 6F | >0.9999 |
| TgF344-AD 9M vs. 9F | >0.9999 |
| TgF344-AD 12M v. 12F | >0.9999 |
| **Aβ40** | |
| F344 3M v. 3F | >0.9999 |
| F344 6M v. 6F | 0.8370 |
| F344 12M v. 12F | 0.3586 |

***Location Novelty Recognition Task***

The Location Novelty Recognition Test (LNRT) was performed using a separate group of unimplanted animals as internal control experiment only for the purpose of confirming observations from previous reports indicating α5IA enhances memory function. These data were not relied up to arrive at any conclusion based on *in vivo* electrophysiological studies of the neural correlates of memory function. The LNRT is a simple test of memory function that relies on the rat's innate exploratory behaviors to assess the ability of the animal to recognize that an object is in new location without having to employ any rules or behavioral reinforces. In the current study, animals were acclimated to the same familiar environment used in the *in vivo* electrophysiological studies. During the training trial two copies of the same object were placed in two different quadrants of the familiar environment and the animal was allowed to explore these objects for 10 minutes. Following a two-hour delay, one of the two objects from the training trial was moved in a novel location in a previously vacant quadrant and the animal was reintroduced to this environment and once again allowed to explore the objects for 10 minutes. The preference for novelty is revealed by the tendency of the animal to spend more time exploring the displaced object. Two identical objects were used in a given vehicle drug training and test session but, different objects were used for each rat to ensure counter balancing for differences in the complexity of the object. All objects were tested prior to the beginning of these experiments and found to be devoid of bias – rats spent equal time with each of the objects used. The environment floor and objects were cleaned with 30% ethanol between recordings to remove olfactory cues. A location discrimination index was calculated for each trial as previously described (*31, 32*):

$$Location\ Index = \frac{(T_{NOV}\ x\ 100)}{(T_{NOV} + T_{FAM})}$$

where, $T_{NOV}$ is the time spent exploring the displaced object and $T_{FAM}$ is the time spent exploring the non-displaced object. A one-sample *t*-test was used to determine whether the location index was different from chance level performance (50%) and the performance of each rat in either treatment group was directly compared using a paired, one-tailed *t*-test with a probability level of $\alpha < 0.05$ considered to be statistically significant.

***Validation of Objects used for the LNRT***: Each type of object was evaluated for intrinsic bias by comparing the time spent exploring the type of object with the other objects using the one-way Wilcoxon Test **(Table S4 and Fig S8).** Object 1 (deodorant bottle); Object 2 (bubbles bottle); Object 3 (salt shaker); Object 4 (doll). All objects were thoroughly cleaned, and filled with lead pellets to anchor them in place during trials. The results of a Kruskal-Wallis test comparing the time spent exploring each of the four different types of objects with each other was not significant (p = 0.21) ensuring that rats spent a similar amount of time interacting with each. None of the objects included in these studies had a negative or positive intrinsic value. Animals showed no aversion to any of the objects used in these studies.

**Table S4: Results of One-Way Wilcoxon Test for Object Validation and Evaluation for Intrinsic Bias**

| **One-Way Wilcoxon Test** | **Object 1** | **Object 2** | **Object 3** | **Object 4** | **All Objects** |
|---|---|---|---|---|---|
| Theoretical median | 50.00 | 50.00 | 50.00 | 50.00 | 50.00 |
| Actual median | 35.80 | 55.66 | 39.99 | 82.04 | 39.15 |
| Number of values | 12 | 6 | 4 | 6 | 34 |
| Sum of signed ranks (W) | -14.00 | 9.000 | -4.000 | 15.00 | -165.0 |
| Sum of positive ranks | 32.00 | 15.00 | 3.000 | 18.00 | 215.0 |
| Sum of negative ranks | -46.00 | -6.000 | -7.000 | -3.000 | -380.0 |
| **P value (two tailed)** | **0.6221** | **0.4375** | **0.6250** | **0.1562** | **0.1629** |
| Exact or estimate? | Exact | Exact | Exact | Exact | Exact |
| **P value summary** | **ns** | **ns** | **ns** | **ns** | **ns** |
| **Significant (alpha=0.05)?** | **No** | **No** | **No** | **No** | **No** |

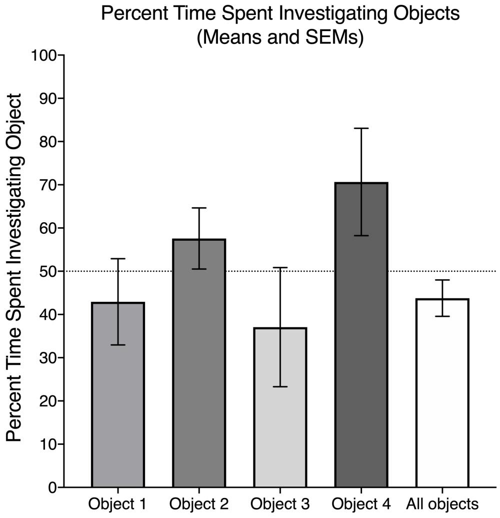


**Fig S8:** Histogram showing Means and SEMs for percent time spent investigating the different types of objects as compare to chance.

##### *Statistical Analysis of Data*

The unit of analysis for each comparison was selected based on the dependent variable interest. The unit of analysis of the escalating dose, place cell remapping activity and cell by cell comparisons was the single pyramidal cell or interneuron. The unit of analysis for LFPs power at each frequency was an FFT point. The unit of analysis for the location novelty recognition behavioral testing was an animal. Data were analyzed for normality prior to statistical analysis using the Kolmogorov-Smirnov test. Normally distributed data was analyzed for within subject effects using a parametric paired subject t tests and repeated measures ANOVAs with Bonferroni corrections for multiple comparisons. The non-parametric Wilcoxon Signed Rank test, Kruskal-Wallis test and Friedman's ANOVA were used for analysis of data that was not normally distributed. Planned comparisons were performed based on dose-dependent *a priori* expected effects of drug-mediated disinhibition on mean and peak firing rates, spatial information content (SIC), and LFP power all of which have previously been shown to be sensitive to drug-induced changes in inhibition and excitation. All results are expressed as the mean ± SEMs with an alpha $p < 0.05$ significance level unless otherwise indicated. The alpha level was adjusted accordingly for the number of multiple comparisons being made in a repeated measures analysis. All statistical analyses were performed using the SPSS Statistics Program for Macintosh, Version 20.0 (IBM Corporation, Armonk, NY, USA). Figures were generated using GraphPad Prism (GraphPad Software, La Jolla California USA).

#### SUPPLEMENTAL RESULTS

##### *Analysis of Mean Firing Rates of LE Rat Place Cells in Escalating Dose Model*

The firing rates of LE rat CA1 pyramidal cells were measured while animals explored the familiar environment following administration of vehicle and escalating doses of $\alpha$5IA. The average percent change in mean firing rate ranged from 23 to 71% with an average change of 38% following administration of the 1.0 mg/kg dose of $\alpha$5IA (**Table S5**). The effect of vehicle versus $\alpha$5IA on the activity of individual cells in the scatter plots and frequency distributions (**Fig S9).**

A B C

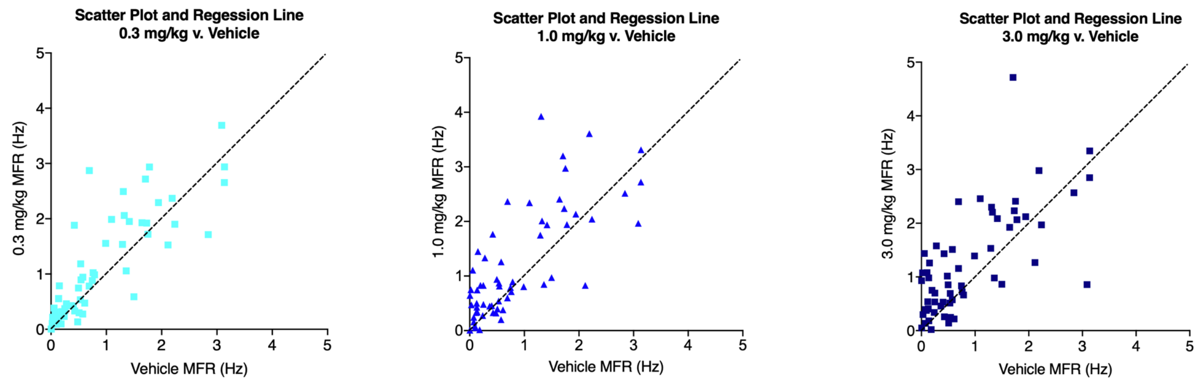


**Fig. S9. Scatter Plots for Mean Firing Rates in Escalating Dose Model. A)** Scatter plot for vehicle fit curve line versus 0.3 mg/kg dose of $\alpha$5IA. **B)** Scatter plot for vehicle versus 1.0 mg/kg. **C)** Scatter plot for vehicle versus 3.0 mg/kg.

**Table S5: Mean Firing Rates of Individual LE Rats Escalating Dose Model**

| Rat # | Vehicle | 0.3 mg/kg | 1.0 mg/kg | 3.0 mg/kg | % Change (1.0 mg/kg) |
|---|---|---|---|---|---|
| 204761 | 1.08665905 | 1.22439015 | **1.34093528** | 1.48529932 | 23% |
| 206440 | 0.74717198 | 0.95881221 | **1.27890491** | 1.17652844 | 71% |
| 207232 | 0.96529718 | 1.40989799 | **1.28803643** | 1.31341157 | 33% |
| 123408 | 0.46385454 | 0.60343560 | **0.61672310** | 0.58985254 | 33% |
| Average | 0.81574568 | 1.04913398 | **1.13114993** | 1.14127296 | **38%** |

##### *Supplemental Results from Remapping Model Analyses*

Oral $\alpha$5IA administration significantly increased the mean, peak and in-field mean firing rates (MFR) of CA1 place cells in this remapping model in the LE Rats (Rats n= 3; Place cells n = 115). Planned comparisons indicated that $\alpha$5IA administration significantly increased MFRs in the familiar environment ($p < 0.01$) **(Fig. S10-A and S11; Tables S6, S7 and S8).** The increase MFR of CA1 place cells seen in the LE rats is consistent with that observed in the F344 rats **(see Fig. 2 in manuscript).** Planned comparisons also revealed significant increases in MFR on exposure to environmental novelty in the vehicle conditions indicating an effect of environment on mean firing rates as observed in F344 rats **(Fig. S10-A; Table S8)**. These effects of drug and environment were significant after adjusting the alpha for multiple comparisons. The interaction between drug and environment was not significant on this parameter. Peak firing rates (PFR) also increased significantly in the familiar environment following $\alpha$5IA administration p<0.01). **(Fig. S10-B; Table S8)** Planned comparisons also revealed significant increases in PFR during exposure to environmental novelty in the vehicle (p<0.05). Although this was not significant after controlling for multiple comparisons it is important to note that this control experiment was not influenced by subsequent experiments in the drug conditions **(Fig. S10-B; Table S8)**. The interaction between drug and environment was not significant on this parameter. Significant in-field MFRs changes are also seen in this model. **(Fig. S10-C; Table S8)** Planned comparisons revealed a significant in-field MFR increase in the familiar environment following $\alpha$5IA administration. Planned comparisons indicated that in-field MFRs also increased significantly in the novel environment under both the vehicle indicating an effect of environment on this parameter. The effect of novelty on this parameter in the drug condition was significant after controlling for multiple comparisons. The interaction between drug and environment was not significant on this parameter.

Analysis of variance for SIC per spike revealed significant effects. **(Fig. S10-D; Table S8)** Planned comparisons demonstrated significant decreases in SIC in the novel environment in the vehicle and drug treatment conditions. The interaction between drug and environment was not significant on this parameter.

Significant changes in place field area (PFA) were revealed by ANOVA as well. **(Fig. S10-E; Table S8)** PFA increased with environmental novelty from 366cm$^2$ to 407cm$^2$ (11%) and 368cm$^2$ to 412 cm$^2$ (12%) in the vehicle and drug conditions respectively but, although there was a strong trend (p = 0.07) observed in vehicle condition the increase in PFA only reached statistical significance (p = 0.001) in the drug condition. The interaction between drug and environment was not significant on this parameter.

Analysis of variance also revealed significant differences in place field spatial correlations in the vehicle condition. **(Fig. S10-F; Table S8)** Pairwise comparisons indicated spatial correlations between first familiar and each of the two novels were significantly lower than the correlations between the two familiars indicating remapping occurred during both exposures to the novel in the vehicle condition. This effect remained significant when controlling for multiple comparisons. Significant differences in spatial correlations were also revealed by ANOVA in the drug condition. Pairwise comparisons indicated spatial correlations between first familiar and each of the two novels were significantly lower than the correlations between the two familiars indicating remapping occurred during both exposures to the novel in the drug condition as well. Comparisons of spatial correlations across treatment conditions revealed no differences in the correlation between the two familiar environments before or after controlling for multiple comparisons indicating no effect on drug on the stability of well-established highly correlated place fields in this model (*37*). The interaction between drug and environment was not significant for this parameter.

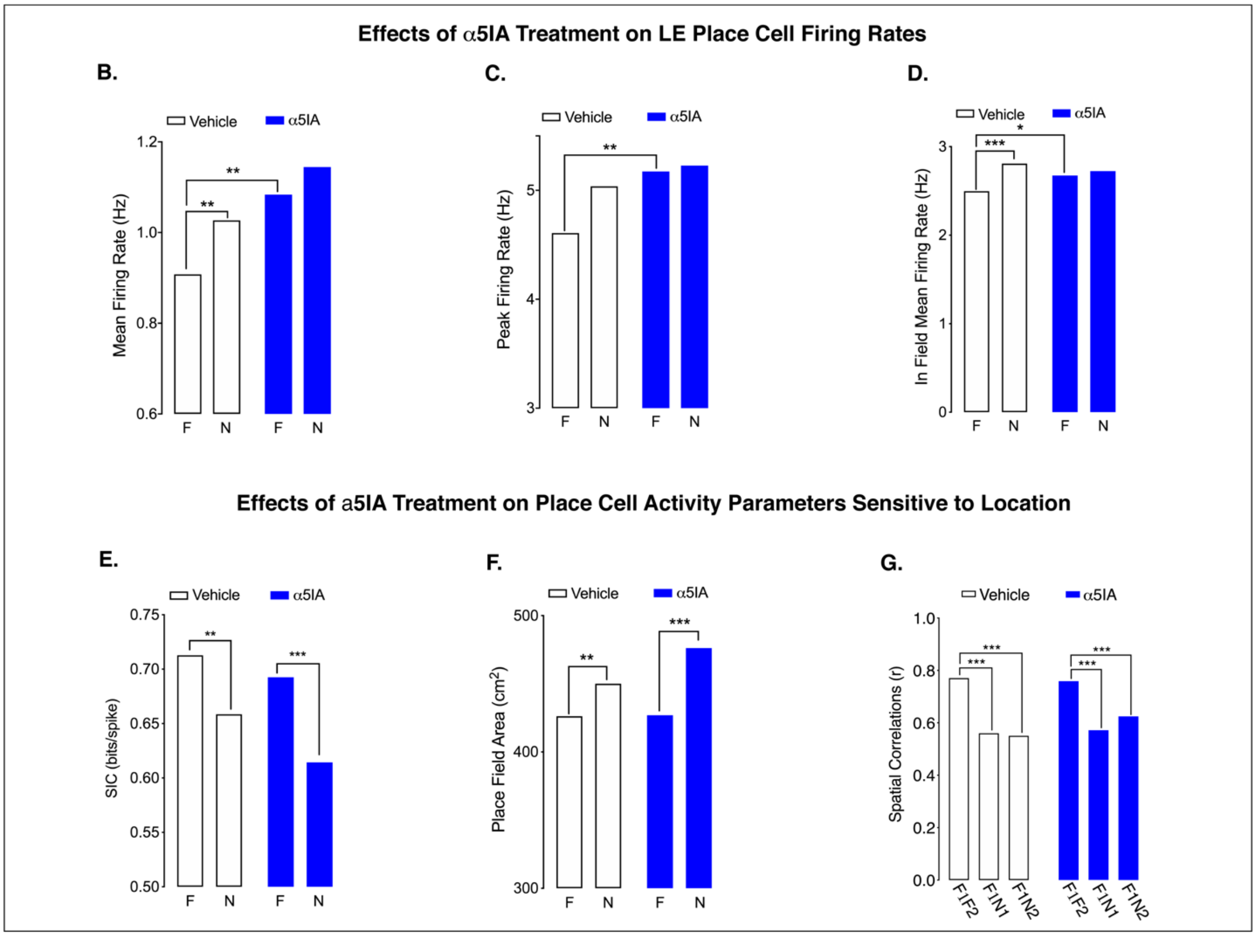


**Fig. S10. Results of remapping model experiments designed to assess the within-subject effects of oral α5IA administration on CA1 place cell responses to environmental novelty in adult male LE rats. A)** Environment-dependent remapping model. **A)** Histogram shows significant effect of α5IA (1.0 mg/kg) on MFR of CA1 place cells in a familiar environment and significant effects of environmental novelty on MFR. **B)** Significant effect of α5IA on PFR of place cells in a familiar environment. **C)** Effect of α5IA on in-field MFR of CA1 place cells in a familiar environment and significant effects of environmental novelty on in-field MFR. **D)** SIC significantly decreases during exposure to environmental novelty. **E)** Planned comparisons indicate that PFA significantly increases following administration of α5IA but ANOVA reveals no interaction between environment and treatment on this parameter. **F)** Environmental novelty significantly reduces place field spatial correlations novelty in both treatment conditions. Rats n = 3. Place cells n = 115. Significant at $*p<0.01$ unless otherwise indicated.

**Table S6: Means and SEMs for Remapping Data from LE Rats**

| **Parameter** | **Mean** | **SEM** |
|---|---|---|
| MFR Familiar Vehicle | 0.91 | ± 0.062 |
| MFR Novel Vehicle | 1.03 | ± 0.073 |
| MFR Familiar α5IA | 1.08 | ± 0.083 |
| MFR Novel α5IA | 1.14 | ± 0.093 |
| PFR Familiar Vehicle | 4.61 | ± 0.344 |
| PFR Novel Vehicle | 5.04 | ± 0.359 |
| PFR Familiar α5IA | 5.17 | ± 0.395 |
| PFR Novel α5IA | 5.23 | ± 0.338 |

| | | |
|---|---|---|
| In Field MFR Familiar Vehicle | 2.50 | ± 0.169 |
| In Field MFR Novel Vehicle | 2.67 | ± 0.150 |
| In Field MFR Familiar α5IA | 2.74 | ± 0.184 |
| In Field MFR Novel α5IA | 2.80 | ± 0.166 |
| SIC Familiar Vehicle | 0.71 | ± 0.046 |
| SIC Novel Vehicle | 0.69 | ± 0.044 |
| SIC Familiar α5IA | 0.66 | ± 0.050 |
| SIC Novel α5IA | 0.61 | ± 0.041 |
| Area Familiar Vehicle | 426 | ± 14.81 |
| Area Novel Vehicle | 450 | ± 18.50 |
| Area Familiar α5IA | 427 | ± 15.25 |
| Area Novel α5IA | 476 | ± 18.13 |
| Correlation F1F4 Vehicle | 0.77 | ± 0.013 |
| Correlation F1N1 Vehicle | 0.56 | ± 0.027 |
| Correlation F1N2 Vehicle | 0.55 | ± 0.027 |
| Correlation F1F4 α5IA | 0.75 | ± 0.014 |
| Correlation F1N1 α5IA | 0.57 | ± 0.023 |
| Correlation NF1N2 α5IA | 0.62 | ± 0.023 |

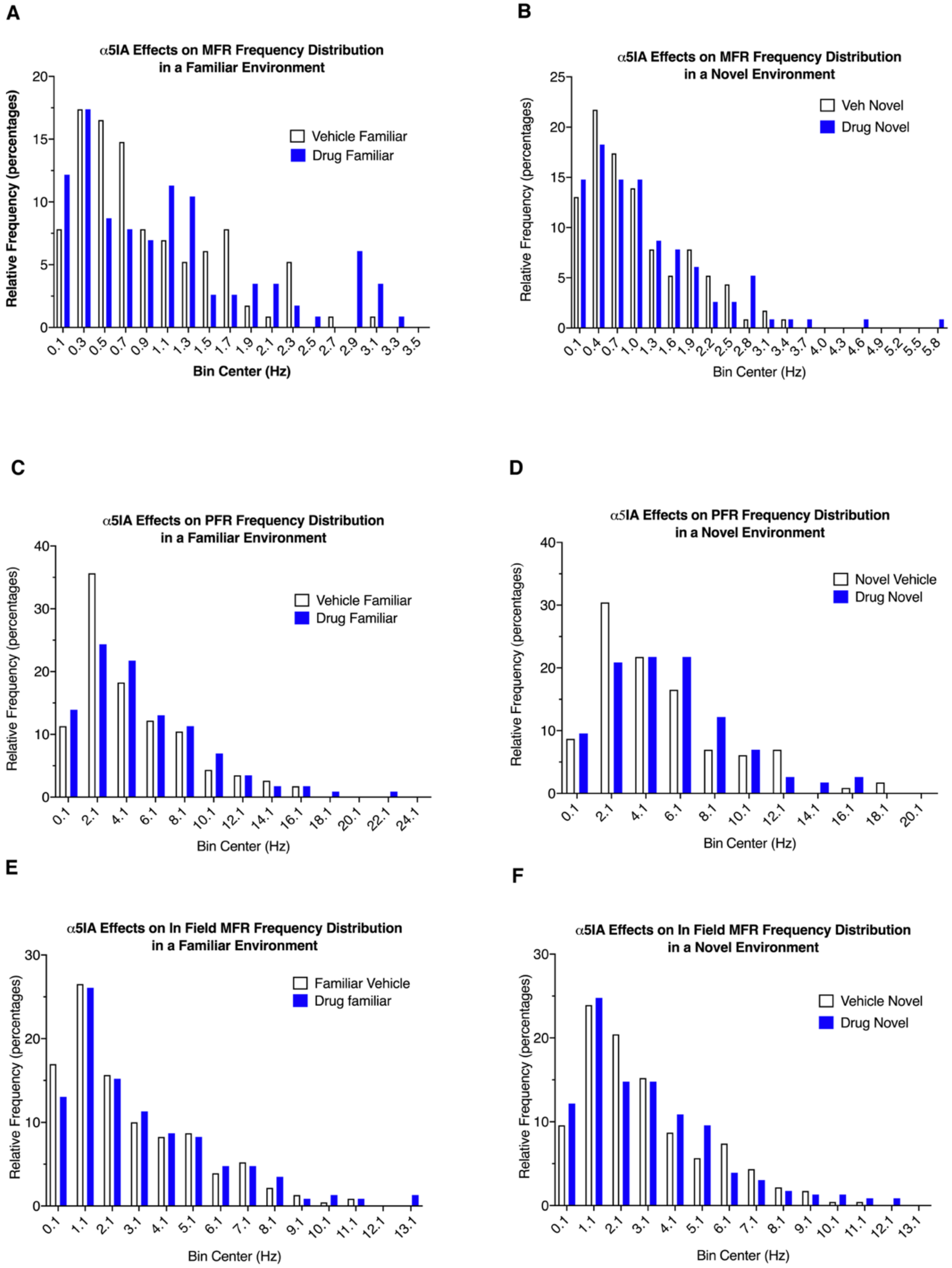


**Fig S11. Firing Rate Frequency Distributions Reveal Effects of α5IA on Mean and Peak Firing Rates in the Familiar and Novel Environments in LE rats. A)** The mean firing rate frequency distribution in the familiar environment reveals a shift in the distribution with a remarkable increase in the percentage of cells with mean firing rates $\geq 1.9$

Hz following α5IA administration. **B)** Mean firing rate frequency distributions in the novel environment also reveals a shift in the frequency distribution with an increase in the percentage of cells with mean firing rates ≥ 2.8 Hz following α5IA administration. **C)** Peak firing rate frequency distributions in the familiar environment also reveals a shift in the frequency distribution with an increase in the percentage of cells with peak firing rates ≥ 4.1 Hz following α5IA administration. **D)** Peak firing rate frequency distributions in the novel environment also reveals a shift in the frequency distribution with an increase in the percentage of cells with peak firing rates ≥ 6.1 Hz following α5IA administration. **E)** In-field mean firing rate frequency distributions in the familiar environment also reveals a shift in the frequency distribution with an increase in the percentage of cells with peak firing rates ≥ 8.1 Hz following α5IA administration. **F)** In-field mean firing rate frequency distributions in the novel environment also reveals a shift in the frequency distribution with an increase in the percentage of cells with peak firing rates ≥ 10.1 Hz following α5IA.

**Table S7. Summary of Significant Effects of Oral α5IA on CA1 Place Cells in LE Rats**

| **Parameter** | **Drug Familiar** | **Drug Novel** | **Response to Novelty** | **Drug/Env Interaction** |
|---|---|---|---|---|
| Mean FR (MFR) | Yes ↑ | No | Yes ↓ | No |
| Peak FR (PFR) | Yes ↑ | No | No | Trend |
| In Field MFR | Yes ↑ | No | Yes ↓ | No |
| SIC | No | No | Trend | No |
| PF Area (PFA) | No | No | No | No |
| Spatial Correlations (SC) | No | No | Trend for F1:N1 only | No |

Significant effects indicated by yes or no at $p < 0.05$. Trends indicated at $p < 0.1$

**Table S8. Results of Statistical Analysis of Place Cell and Ripple Response to Oral α5IA in LE Rats**

| **Figure #** | **Panel** | **Test statistic (DF) = p value (Dependent Variable)** | **Significance of Pairwise Comparisons** |
|---|---|---|---|
| Fig. 1. | B | *Friedman's ANOVA (F (3, 86) = 7.24; p = 0.003) (Escalating Dose Mean Firing Rate)* | Veh v. 0.3mg/kg; p = 0.03*<br>Veh v. 1.0mg/kg; p = 0.0002***<br>Veh v. 2.0mg/kg; p = 0.001** |
| Fig. 2. | B | *ANOVA (F (3,115) = 4.64; p = 0.002) (Mean Firing Rate)* | $Fam_{Veh}$ v. $Nov_{Veh;}$ p = 0.01**<br>$Fam_{Veh}$ v. $Fam_{Drg}$; p = 0.002** |
| | C | *Friedman's ($\chi^2$(3, 115) = 21.91; p < 0.001) (Peak Mean Firing Rate)* | $Fam_{Veh}$ v. $Nov_{Veh;}$ p = 0.05†<br>$Fam_{Veh}$ v. $Fam_{Drg}$; p = 0.002** |
| | D | *Friedman's ($\chi^2$(3, 115) = 17.6; p < 0.001) (In-Field Mean Firing Rate)* | $Fam_{Veh}$ v. $Nov_{Veh;}$ p < 0.001***<br>$Fam_{Veh}$ v. $Fam_{Drg}$; p = 0.013* |
| | E | *Friedman's ($\chi^2$(3, 115) = 25.9; p < 0.001) (Spatial Information Content)* | $Fam_{Veh}$ v. $Nov_{Veh;}$ p < 0.006**<br>$Fam_{Drg}$ v. $Nov_{Drg;}$ p < 0.001***<br>$Fam_{Veh}$ v. $Fam_{Drg}$; p = 0.959<br>$Nov_{Veh}$ v. $Nov_{Drg}$; p = 0.645 |
| | F | *Friedman's ($\chi^2$ (3, 115) = 20.3; p < 0.001) (Place Field Area)* | $Fam_{Veh}$ v. $Nov_{Veh;}$ p < 0.005**<br>$Fam_{Drg}$ v. $Nov_{Drg;}$ p < 0.001*** |

| | | | |
|---|---|---|---|
| | G | *Friedman's ($\chi^2$(2, 77) = 63.3.0; p < 0.001)*<br>*(Spatial Correlations)* | $Fam_{Veh}$ v. $Fam_{Drg}$; p = 0.867<br>$Nov_{Veh}$ v. $Novrg_{D}$; p = 0.767<br><br>$Fam_{Veh}$ v. $Nov_{Veh1;}$ p < 0.001***<br>$Fam_{Veh}$ v. $Nov_{Veh2;}$ p < 0.001***<br>$Fam_{Drg}$ v. $Nov_{Drg1;}$ p < 0.001***<br>$Fam_{Drg}$ v. $Nov_{Drg1;}$ p < 0.001***<br>$Fam_{Veh}$ v. $Fam_{Drg}$; p = 0.86 |
| Fig. S3. | B | *ANOVA (F (3, 86) = 7.24; p = 0.003)*<br>*(Escalating Dose Mean Firing Rate)* | Veh v. 1.0mg/kg; p = 0.003**<br>Veh v. 2.0mg/kg; p = 0.02*<br>1.0mg v. kg/3.0mg/kg; p > 0.999 |
| Fig. 4. | E | *Friedman's ($\chi^2$ (4, 125) = 361; p < 0.001)*<br>*(Mean Power Spectral Density)* | $PSD_{Veh}$ v. $PSD_{0.3mg/kg}$; p < 0.001***<br>$PSD_{Veh}$ v. $PSD_{1.0mg/kg}$; p < 0.001***<br>$PSD_{Veh}$ v. $PSD_{3.0mg/kg}$; p < 0.001*** |
| | F | *Friedman's ($\chi^2$ (1, 3) = 12; p = 0.007)*<br>*(Peak Power Spectral Density)5* | $PSD_{Veh}$ v. $PSD_{0.3mg/kg}$; p = 0.02†<br>$PSD_{Veh}$ v. $PSD_{1.0mg/kg}$; p = 0.007**<br>$PSD_{Veh}$ v. $PSD_{3.0mg/kg}$; p = 0.273 |
| | H | *Wilcoxon* ($\chi^2$ (1, 125) = 9.16; p < 0.001)<br>*(Mean Power Spectral Density no Theta)* | $PSD_{Veh}$ v. $PSD_{D}$; p < 0.001*** |
| | K | *Friedman's ($\chi^2$ (4, 125) = 312; p < 0.001)*<br>*(Mean Power Spectral Density)* | $PSD_{Veh}$ v. $PSD_{0.3mg/kg}$; p > 0.05<br>$PSD_{Veh}$ v. $PSD_{1.0mg/kg}$; p < 0.001***<br>$PSD_{Veh}$ v. $PSD_{2.0mg/kg}$; p < 0.001*** |
| | L | *Friedman's ($\chi^2$ (1, 3) = 13; p = 0.004)*<br>*(Peak Power Spectral Density)* | $PSD_{Veh}$ v. $PSD_{0.3mg/kg}$; p = 0.462<br>$PSD_{Veh}$ v. $PSD_{1.0mg/kg}$; p = 0.01**<br>$PSD_{Veh}$ v. $PSD_{2.0mg/kg}$; p = 0.004** |
| | N | *Wilcoxon ($\chi^2$ (1, 125) = 22; p < 0.001)*<br>*(Mean Power Spectral Density no Theta)* | $PSD_{Veh}$ v. $PSD_{D}$; p < 0.001*** |
| | O | *Friedman's ($\chi^2$ (3, 125) = 43; p < 0.001)*<br>*(Mean Power Spectral Density)* | $PSD_{Veh}$ v. $PSD_{0.3mg/kg}$; p = 0.462<br>$PSD_{Veh}$ v. $PSD_{1.0mg/kg}$; p < 0.001***<br>$PSD_{Veh}$ v. $PSD_{2.0mg/kg}$; p < 0.001*** |
| | Q | *Friedman's* ($\chi^2$ (1, 3) = 11; p = 0.006)<br>*(Peak Power Spectral Density)* | $PSD_{Veh}$ v. $PSD_{0.3mg/kg}$; p = 0.055†<br>$PSD_{Veh}$ v. $PSD_{1.0mg/kg}$; p = 0.17<br>$PSD_{Veh}$ v. $PSD_{2.0mg/kg}$; p = 0.006** |
| | R | *Wilcoxon* ($\chi$ (1, 125) = 18; p < 0.001)<br>*(Mean Power Spectral Density no Theta)* | $PSD_{Veh}$ v. $PSD_{D}$; p < 0.001*** |
| Fig. 5. | B | *Kruskal-Wallis* (H (3, 405) = 15; p < 0.0001)<br>*(Ripple Peak Amplitude)* | $Amp_{Veh}$ v. $Amp_{0.3mg/kg}$; p < 0.001***<br>$Amp_{Veh}$ v. $Amp_{1.0mg/kg}$; p < 0.001***<br>$Amp_{Veh}$ v. $Amp_{2.0mg/kg}$; p < 0.001*** |
| | C | *ANOVA F (3, 86) = 6.13; p = 0.018)* | $Number_{Veh}$ v. $Number_{0.3mg/kg}$; p > 0.724 |

| | | | |
|---|---|---|---|
| | | *(Ripple Counts)* | $Number_{Veh}$ v. $Number_{1.0mg/kg}$; p = 0.022* |
| | D | *Kruskal-Wallis* (H *(3, 2066) = 11.9; p = 0.008)* *(Ripple Duration)* | $Dur_{Veh}$ v. $Dur_{0.3mg/kg}$; p = 0.943<br>$Dur_{Veh}$ v. $Dur_{1.0mg/kg}$; p = 0.003*<br>$Dur_{Veh}$ v. $Dur_{2.0mg/kg}$; p > 0.180 |
| | E | *Kruskal-Wallis ($\chi^2$ (3, 2066); p = 0.124)* *(Ripple Peak Frequency)* | |

Significant effects indicated by: * at p < 0.05; ** at p < 0.01; and, *** at p < 0.001.
Trends not significant after controlling for multiple comparisons indicated by †.

***Supplement Data for Place Cell Firing Analysis in F344 Adult Male Rats***

We successfully recorded data from n = 152 place cells in three F344 adult male rats. All data were analyzed using Friedman's non-parametric repeated measures ANOVA with alpha level adjusted for the number of multiple pairwise comparisons unless otherwise indicated.

Friedman's ANOVA for mean firing rate was significant (F (3, 152) = 51.7; p < 0.001). Planned comparisons indicated that $\alpha$5IA administration significantly increased MFRs by 18.5% in the familiar environment (p < 0.001) **(see Fig. 2A in manuscript).** Planned comparisons also revealed significant increases in MFR on exposure to environmental novelty in both the vehicle (p<0.001) and drug (p=0.007) conditions indicating an effect of environment on mean firing rates. These effects of drug and environment were all significant after adjusting the alpha for multiple comparisons. The interaction between drug and environment was not significant on this parameter.

Peak firing rates (PFR) also increased significantly following $\alpha$5IA administration ($\chi^2$(3, 152) = 43.3; p<0.001). A planned comparison revealed a small but significant (p=0.01) 7% peak firing rate increase in the familiar environment following $\alpha$5IA administration. **(see Fig. 2B in manuscript)** Planned comparisons also revealed significant increases in PFR during exposure to environmental novelty in both the vehicle (p<0.001) and drug (p=0.01) conditions indicating an effect of environment on peak firing rates. The effect of environmental novelty on PFR in the drug condition remained significant after controlling for multiple comparisons. The interaction between drug and environment was not significant on this parameter.

Significant ($\chi^2$(3, 152) = 38.7; p<0.001) in-field MFRs changes are also seen in this model. **(see Fig. 2C in manuscript)** Planned comparisons revealed a significant (p = 0.004) 11% in-field MFR increase in the familiar environment following $\alpha$5IA administration. Planned comparisons indicated that in-field MFRs also increased significantly in the novel environment under both the vehicle (p<0.001) and drug (p=0.01) conditions indicating an effect of environment on this parameter. The effect of novelty on this parameter in the drug condition was not significant after controlling for multiple comparisons. The interaction between drug and environment was not significant on this parameter.

Analysis of variance in CA1 place cell spatial information content (SIC) per spike in this model also revealed significant ($\chi^2$(3, 152) = 46.0; p < 0.001) effects. **(see Fig. 2D in manuscript)** Planned comparisons demonstrated a significant (p<0.01) decreases in SIC in the novel environment in the vehicle and drug treatment conditions of 18% and 10% respectively. The interaction between drug and environment was not significant on this parameter.

Significant ($\chi^2$ (3, 152) = 12.4; p<0.006) changes in place field area (PFA) were revealed by ANOVA as well. **(see Fig. 2E in manuscript)** PFA increased with environmental novelty from 366cm$^2$ to 407cm$^2$ (11%) and 368cm$^2$ to 412 cm$^2$ (12%) in the vehicle and drug conditions respectively but, although there was a strong trend (p = 0.07) observed in vehicle condition the increase in PFA only reached statistical significance (p = 0.001) in the drug condition. The interaction between drug and environment was not significant on this parameter.

Analysis of variance also revealed significant differences in place field spatial correlations in the vehicle condition ($\chi^2$(2, 44) = 28.7; p < 0.001). **(see Fig. 2F in manuscript)** Pairwise comparisons indicated spatial correlations between first familiar and each of the two novels (F1N1: r = 0.5, p < 0.001; F1N2: r = 0.48, p < 0.001) were significantly lower than the correlations between the two familiars (F1F2: r = 0.66) indicating remapping occurred during both exposures to the novel in the vehicle condition. This effect remained significant when controlling for multiple comparisons. Significant ($\chi^2$(4, 44) = 19.4; p < 0.001) differences in spatial correlations were also revealed by ANOVA in the drug condition. Pairwise comparisons indicated spatial correlations between first familiar and each of the two novels (F1N1: r = 0.59, p < 0.002; F1N2: r = 0.56, p < 0.001) were significantly lower than the correlations between the two familiars (F1F2: r = 0.69) indicating remapping occurred during both exposures to the novel in the drug condition as well. Comparisons of spatial correlations across treatment conditions revealed no differences in the correlation between the two familiar environments before (p=0.20) or after controlling for multiple comparisons (p; $r_{veh}$ = 0.66±0.10; $r_{drug}$ = 0.69±0.11) indicating no effect on drug on the stability of well-established highly correlated place fields in this model (*37*). The interaction between drug and environment was not significant for this parameter.

**Table S9. Means and SEMs for Remapping Data from F344s**

| Parameter | Mean (Hz) | SEM (Hz) |
|---|---|---|
| MFR Familiar Vehicle | 0.74 | ± 0.047 |
| MFR Novel Vehicle | 0.90 | ± 0.057 |
| MFR Familiar α5IA | 0.88 | ± 0.053 |
| MFR Novel α5IA | 0.98 | ± 0.059 |
| PFR Familiar Vehicle | 3.80 | ± 0.213 |
| PFR Novel Vehicle | 4.44 | ± 0.240 |
| PFR Familiar α5IA | 5.07 | ± 0.210 |
| PFR Novel α5IA | 4.86 | ± 0.222 |
| In Field MFR Familiar Vehicle | 2.38 | ± 0.130 |
| In Field MFR Novel Vehicle | 2.76 | ± 0.144 |
| In Field MFR Familiar α5IA | 2.64 | ± 0.137 |
| In Field MFR Novel α5IA | 2.93 | ± 0.140 |
| SIC Familiar Vehicle | 0.73 | ± 0.042 |
| SIC Novel Vehicle | 0.59 | ± 0.034 |
| SIC Familiar α5IA | 0.70 | ± 0.036 |
| SIC Novel α5IA | 0.60 | ± 0.033 |
| Area Familiar Vehicle | 366 | ± 13.48 |
| Area Novel Vehicle | 408 | ± 20.25 |
| Area Familiar α5IA | 369 | ± 13.56 |
| Area Novel α5IA | 413 | ± 16.36 |
| Correlation F1F4 Vehicle | 0.66 | ± 0.016 |
| Correlation F1N1 Vehicle | 0.50 | ± 0.025 |

| | | |
|---|---|---|
| Correlation F1N2 Vehicle | 0.48 | $\pm$ 0.028 |
| Correlation F1F4 $\alpha$5IA | 0.69 | $\pm$ 0.016 |
| Correlation F1N1 $\alpha$5IA | 0.60 | $\pm$ 0.022 |
| Correlation NF1N2 $\alpha$5IA | 0.56 | $\pm$ 0.027 |

***Supplemental Analysis for Cell by Cell Effects of Oral $\alpha$5IA on Activity of Individual LE CA1 Pyramidal Cell and Interneurons***

Results of the cell by cell analysis for the percentages of pyramidal neurons increasing versus decreasing MFRs was performed using data from the 4 LE rats in included in this study. This analysis revealed that significantly ($\chi^2$ = 8.1; DF = 2; p<0.0174; 45% (60/134) more of these pyramidal cells increased MFR by $\geq$ 20% in a familiar environment after $\alpha$5IA administration; a finding consistent with the data from the escalating dose-response experiment **(Table S10 and Fig. S12).**

Chi square analysis of interneuron activity indicates a significantly ($\chi^2$ = 22.5; DF = 1; p < 0.0001) greater portion of cells 56% (19/34) increased their MFR by $\geq$ 20% following administration of $\alpha$5IA, while only 18% (6/34) showed a decrease of this magnitude. The average MFR of these interneurons across are all 4 environments following vehicle and $\alpha$5IA administration were 12.2 and 16.2 Hz respectively. The results of a pair subject T-test was also significant (t (33) = 2.08; p = 0.045) demonstrating that $\alpha$5IA administration modulates the activity of interneurons as well as pyramidal cells in the CA1 subregion.

Place cells were also characterized based on the percentage of cells that rate remapped following vehicle versus $\alpha$5IA administration. The number of place cells that rate remapped in the vehicle and drug conditions were 31% (42/134) and 25% (33/134) respectively. The observed percentage of cells showing rate remapping in this model which used different shaped environments in the same room is consistent with previous reports (*S4*). Of the place cells that showed a $\geq$ 20% increase in overall mean firing rate following drug administration, 25% (15/60) also showed rate remapping based on a $\geq$ 50% change in their in-field mean firing rates. A slightly smaller percentage 21% (9/42) of the place cells that showed a decrease in overall mean firing rate with following $\alpha$5IA administration also showed rate remapping in the drug condition. A similar percentage 21% (7/33) of the place cells that did not show a change of this magnitude in overall mean firing rate due to drug still showed rate remapping suggesting rate remapping does not require a drug-induced change in overall mean firing rate. A similar number of these place cells (vehicle = 11; drug = 10) showed global remapping based on a change in their spatial correlations as well as a change in their firing rates in the vehicle control and drug conditions respectively.

**Table S10. Cell by Cell Mean Firing Rate Responses of Pyramidal Cells and Interneurons to 1.0 mg/kg $\alpha$5IA**

| **Neuron type** | **D +** | **D -** | **D0** | **Local Network Changes (% cells with new activity patterns)** |
|---|---|---|---|---|
| Pyramidal Cell (Pyr) | 60 Pyr (+) | 41 Pyr (-) | 33 Pyr (0) | (\|60(+) + 41(-)\|) = 75% (101/134) of pyramidal cells |
| Interneurons (Int) | 19 Int (+) | 6 Int (-) | 9 Int (0) | (\|19(+) + 6(-)\|) = 73% (25/34)of interneurons |
| Total Neurons (Units) | \|79 Tot (+)\| | \|47 Tot (-)\| | 42 Tot (0) | (\|79(+) + 47(-)\|) = 75% (126/168) of total units |

Pyramidal cells and interneurons showing net changes in overall mean firing rate in response to 1.0mg/kg α5IA. A neuron is judged to be excited by drug D+ if MFR(drug)-MFR(veh)/MFR(veh)>,=1.2 MFR(veh); inhibited by drug D- if MFR(drug)-MFR(veh)/MFR(veh)<,=0.8 MFR(veh); and, not responsive to drug D0 if MFR(drug)-MFR(veh)/MFR(veh)<,=1.2 MFR(veh) or >,=0.8 MFR(veh).

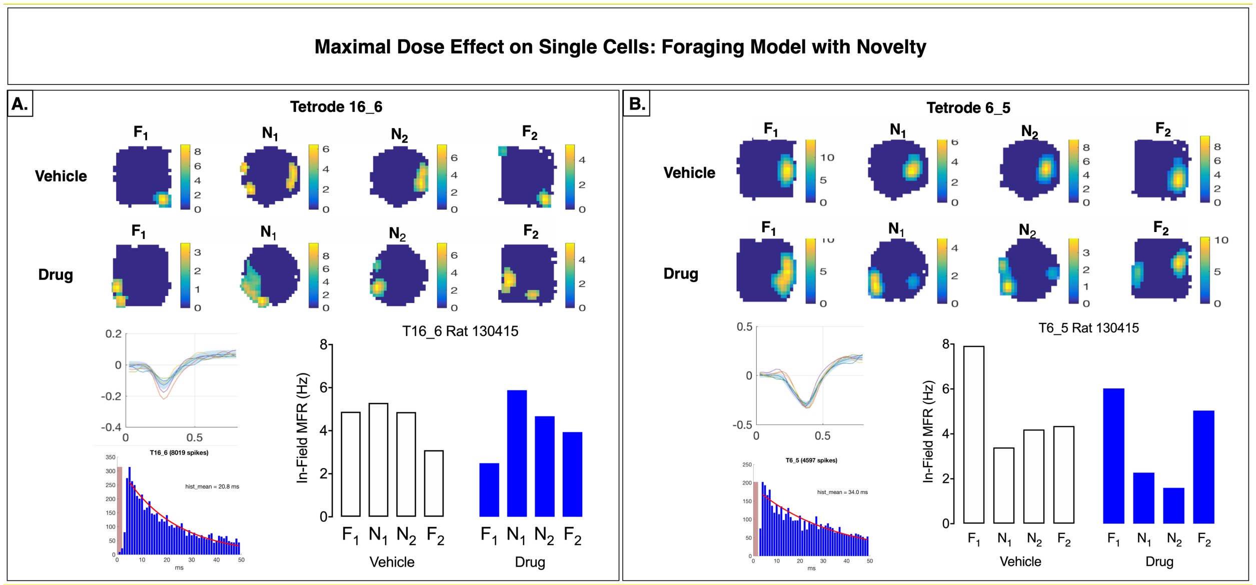


**Fig. S12. Representative place fields showing changes in spatial correlations before and after administration of α5IA in LE Rats**. **A)** CA1 neuron showing overt global remapping of place field heat maps across familiar (F) and novel (N) environments, following oral administration of vehicle (Top: F1-N1-N2-F2), without overt global remapping in the presence of α5IA (Bottom: F1-N1-N2-F2), but cell shows in field firing rate changes with both vehicle and drug (A, bottom middle panel). **B)** A different CA1 neuron showing stable place field heat maps across familiar (F) and novel (N) environments, following oral administration of vehicle (Top: F1-N1-N2-F2), demonstrates global remapping (Bottom: F1-N1-N2-F2) with a change in both location and firing rate (bottom middle panel) following administration of α5IA. The environments for CA1 neuron A and B were: F, square shape, black wood walls and epoxy wood floor, yellow vertical stripes; N1, hexagonal shape, black wood walls, epoxy wood floor, yellow vertical stripes; N2, cylindrical shape, silver metallic walls, black epoxy coated wood floor, yellow vertical stripes.

##### *Supplemental Results for Dose-Dependent Effects of α5IA on Ripples in F344 and TgF344-AD Rats*

To control for systematic error, we performed additional experiments using vehicle only in the wakeful immobility model in two F344 male rats and one TgF344-AD rat age 11 mo. These experiments revealed that repeated exposure to a familiar environment, was consistently associated with a decrease in ripple band power and peak ripple amplitudes in both strains under vehicle control even when the number of ripple events increased **(Fig S13 and S14; Tables S11 and S12).**

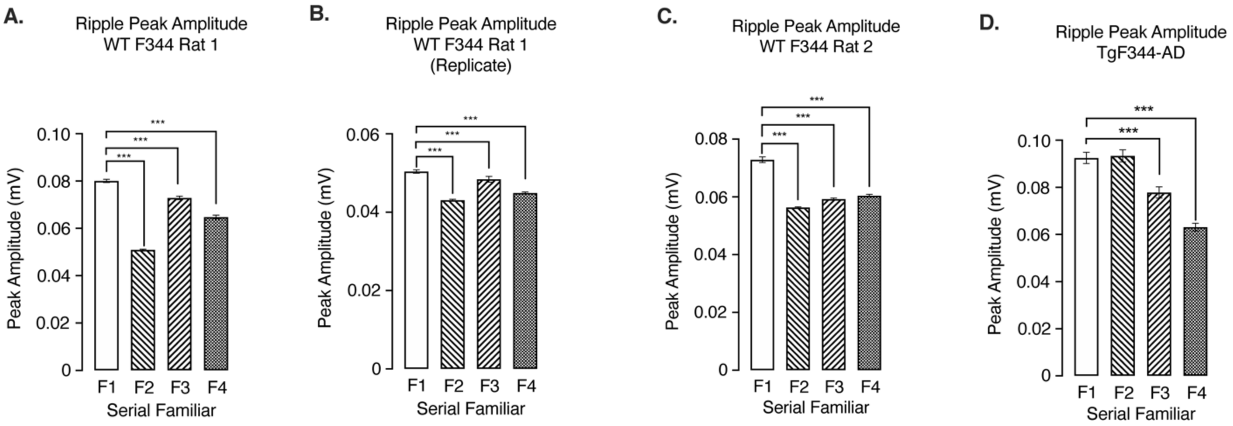


**Fig S13: Wakeful immobility model vehicle control experiments. A** Repeated exposure to a familiar environment is associated with significant decreases in peak ripple amplitude for subject 9. **B)** Results of replicate experiment in this same animal. **C)** Peak ripple amplitudes for subject 10. **D)** Peak ripple amplitudes for TgF344-AD (TgF344-AD subject 8). Significant effects indicated by: *** at p < 0.001.

**Table S11. F344 Vehicle Control Experiments Total Number of Ripples Per Session**

| **Subject (Number)** | **Vehicle** | **Vehicle** | **Vehicle** | **Vehicle** |
|---|---|---|---|---|
| **Subject 1 (F344 Rat #9)** | 502 | 579 | 430 | 409 |
| **Subject 1 (F344 Rat #9)** | 463 | 605 | 435 | 608 |
| **Subject 2 (F344 Rat #10)** | 320 | 648 | 558 | 508 |
| **Subject 3 (Tg344-AD Rat #8)*** | 579 | 564 | 472 | 508 |
| **Average for all F344s** | 428 | 611 | 474 | 417 |
| **Percent change F344s** | N/A | 150% | 118% | 124% |
| **Statistical significance** | N/A | p = 0.094 | p = 0.999 | P = 0.999 |

Total number ripple events per 10-minute recording session normalized by periods of immobility.
* See Table S12

**Table S12. F344s and TgF344-AD Vehicle Control Experiments Total Number of Ripples Per Session**

| **Subjects** | **Vehicle** | **Vehicle** | **Vehicle** | **Vehicle** |
|---|---|---|---|---|
| Average for all subjects | 466 | 599 | 474 | 508 |
| Percent change all subjects | N/A | 129% | 102% | 109% |
| Statistical significance | N/A | p = 0.112 | p = 0.999 | P = 0.999 |

Total number ripple events per 10-minute recording session normalized by periods of immobility.

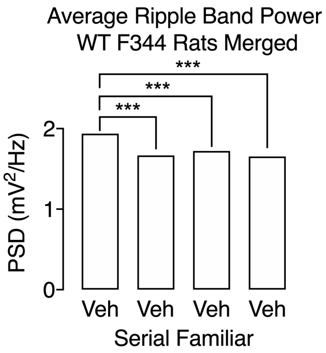


**Fig S14: Average Ripple Band Power F344 Vehicle Control Condition**. Within subject ANOVA reveals reduction in average ripple band power (140 to 200 Hz) in 11 mo old F344 male rats. Significant effects indicated by: *** at $p < 0.001$.

In addition, administration of single bolus doses of 1.0 mg/kg and 2.0 mg/kg of $\alpha$5IA immediately after vehicle to two of the F344 rats used in the escalating dose experiments and one of the rats used in the vehicle control experiments demonstrated that the increase in ripple band power induced by $\alpha$5IA administration are seen under these bolus dosing conditions as well. (**Fig. S15**)

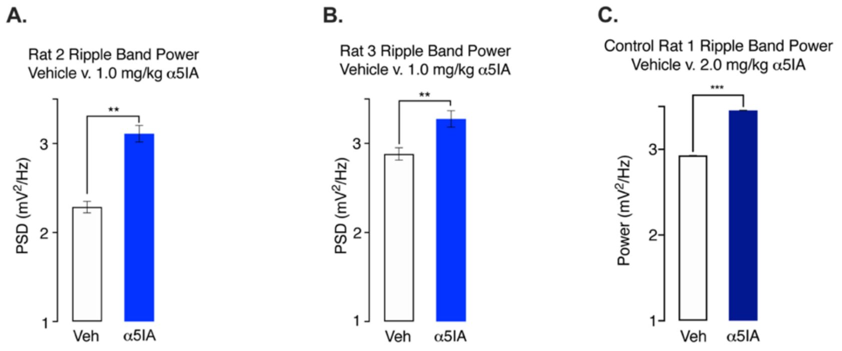


**Fig. S15. Increase in Ripple Band Power induced by single bolus dose of $\alpha$5IA (1.0mg/kg). A)** Single bolus dose of $\alpha$5IA (1.0 mg/kg) in F344 subject 2. **B)** Single bolus dose of $\alpha$5IA (1.0 mg/kg) in F344 subject 3. **C)** Single bolus dose of $\alpha$5IA (2.0 mg/kg) in F344 subject 1. All PSD values are x $10^{-6}$. Significant effects indicated by: ** at $p < 0.01$; and, *** at $p < 0.001$.

***Supplemental Results for Effects of α5IA on Ripple Band Power in Wild Type Rats***

**Table S13: 95% Confidence Intervals for α5IA Escalating Dose PSD Power in Ripple Band of Wildtyp Rats**

| **Rat #1 (15 mo LE)** | **Vehicle** | **0.3 mg/kg** | **1.0 mg/kg** | **2.0 mg/kg** |
|---|---|---|---|---|
| Lower 95% CI | 3.312e-006 | 3.974e-006 | 4.605e-006 | 4.000e-006 |
| Upper 95% CI | 3.493e-006 | 4.217e-006 | 4.935e-006 | 4.240e-006 |
| **Rat #2 (9 mo F344)** | | | | |
| Lower 95% CI | 2.513e-006 | 2.661e-006 | 3.880e-006 | 3.732e-006 |
| Upper 95% CI | 2.745e-006 | 2.932e-006 | 4.322e-006 | 4.199e-006 |
| **Rat #3 (16 mo F344)** | | | | |
| Lower 95% CI | 2.237e-006 | 2.366e-006 | 2.514e-006 | 2.551-006 |
| Upper 95% CI | 2.330e-006 | 2.480e-006 | 2.638e-006 | 2.716e-006 |
| **Rat #4 (12 mo F344)** | | | | |
| Lower 95% CI | 2.668e-006 | 2.668e-006 | 3.234e-006 | 2.723e-006 |
| Upper 95% CI | 2.935e-006 | 2.925e-006 | 3.670e-006 | 3.030e-006 |

1. *Effects of α5IA on Total Number of Ripple Events per Session in Wild Type Rats*

In all four wild type animals tested (one LE and three F344) and all three TgF344-AD rats, treatment with oral α5IA increased the total number of ripples per session. Kruskal-Wallis ANOVA with planned comparisons revealed a significant increase in ripples per session of 49% ($p = 0.022$) following administration of the 1.0 mg/kg oral dose of α5IA in wild type *rats* (**Table S14**). This increase remained significant after controlling for multiple comparisons with an adjusted alpha level of 0.025. No significant effect of α5IA administration on average number of ripples per session is seen in the TgF344-AD rats (**Table S15**).

**Table S14. Oral α5IA Treatment Increases Total Number of Ripple events Per Session**

| | | **Oral Dose of α5IA** | | |
|---|---|---|---|---|
| **Subject Number** | **Vehicle** | **0.3 mg/kg** | **1.0 mg/kg** | ***2.0 mg/kg** |
| **Subject 1 (#LE-327396)** | 212 | 336 | 346 | 345 |
| **Subject 2 (#F344-130)** | 178 | 178 | 245 | 265 |
| **Subject 3 (#F344-600040)** | 123 | 191 | 227 | 224 |
| **Subject 4 (#F344-113)** | 191 | 189 | 236 | 204 |
| **Average for all WT subjects** | 176 | 223.5 | 263.5 | 259.5 |
| **Change from vehicle baseline** | N/A | 27% | ***50%*** | 47% |
| **Statistical significance** | N/A | p = 0.724 | ***p = 0.022*** | N/A† |

Total number ripple events per 10-minute recording session normalized by periods of immobility
*Highest dose administered to this LE male animal was 3.0 mg/kg. †Statistical significance not calculated due to dosing differences. Significance indicated at alpha = 0.025.

**Table S15. Oral α5IA Treatment Increases Total Number of Ripple events Per Session TgF344-AD Rats**

| | | **Oral Dose of α5IA** | | |
|---|---|---|---|---|
| **Subject Number** | **Vehicle** | **0.3 mg/kg** | **1.0 mg/kg** | ***2.0 mg/kg** |
| **TgF344-AD Subject 1 (#128)** | 120 | 127 | 142 | 119 |
| **TgF344-AD Subject 2 (#126)** | 87 | 107 | 140 | 144 |
| **TgF344-AD Subject 3 (#115)** | 223 | 218 | 242 | 248 |
| **Average for all TgF344-AD subjects** | 143 | 151 | 175 | 170 |

| | | | | |
|---|---|---|---|---|
| **Change from vehicle baseline** | N/A | 6% | *22%* | 19% |
| **Statistical significance** | N/A | p = 0.999 | p = 0.924 | p = 0.999 |

Total number ripple events per 10-minute recording session normalized by periods of immobility
Significance indicated at alpha = 0.025.

*2. Effects of $\alpha$5IA on Peak Amplitude of Ripple in Wild Type Rats*

Non-parametric analysis of variance (Kruskal-Wallis ANOVA test) of peak ripple amplitudes revealed significant drug-induced increases in all four wild type animals tested (**Fig. S16**). For subject 1 (LE Rat: #327396), the result was significant ($\chi^2$ (3, 104) = 24.8; p < 0.0001), revealing dose-dependent increase in average peak amplitude of ripple events of 18% (p < 0.0001) and 9% (p < 0.0001) following administration of the 1.0 and 2.0 mg/kg doses of $\alpha$5IA respectively. (**Fig. S16-A**). For subject 2 (F344 Rat: age 9 mo; #130), the ANOVA revealed significant ($\chi^2$ (3, 133) = 72.1; p < 0.0001) dose-dependent increases in average peak amplitude of ripple events of 17% (p < 0.0001), 28% (p < 0.0001) and 34% (p < 0.0001) following administration of the 0.3, 1.0 and 2.0 mg/kg doses of $\alpha$5IA respectively in this rat (**Fig. S16-B**). The ANOVA of peak ripple amplitude data from Subject 3 (F344 Rat: #600040) also revealed significant ($\chi^2$ (3, 104) = 26.0; p < 0.0001) dose-dependent increases in average peak amplitude of 14% (p < 0.0002) and 22% (p < 0.0001) following administration of the 1.0 and 2.0 mg/kg doses of $\alpha$5IA respectively (**Fig. S16-C**). ANOVA of the ripple peak amplitudes from Subject 4 (F344 Rat: #113) was significant ($\chi^2$ (3, 160) = 85.2; p < 0.0001) as well, revealing dose dependent increases of 15% (p < 0.0001), 31% (p < 0.0001) and 20% (p < 0.0001) following administration of the 0.3, 1.0 and 2.0 mg/kg doses of $\alpha$5IA respectively in this animal (**Fig. S16-D**). ANOVA of the merged peak amplitude data from all three F344 rats was also significant ($\chi^2$ (3, 405) = 150; p < 0.0001). Dose-dependent increases in average peak amplitude of ripple events of 11% (p < 0.0007), 26% (p < 0.0001) and 31% (p < 0.0001) are seen following administration of the 0.3, 1.0 and 2.0 mg/kg doses of $\alpha$5IA respectively (**Fig. S16-E**). Analysis of the merged peak amplitude data from all four rats was also significant ($\chi^2$ (2, 509) = 41; p < 0.0001). Planned comparisons revealed dose-dependent increases in average ripple peak amplitude of 10% (p < 0.033) and 24% (p < 0.0001) following administration of the 0.3 and 1.0 mg/kg doses respectively (**Fig. S16-F**). However, the effect on peak amplitude observed at 0.3 mg/kg dose was not significant when controlling for multiple comparisons.

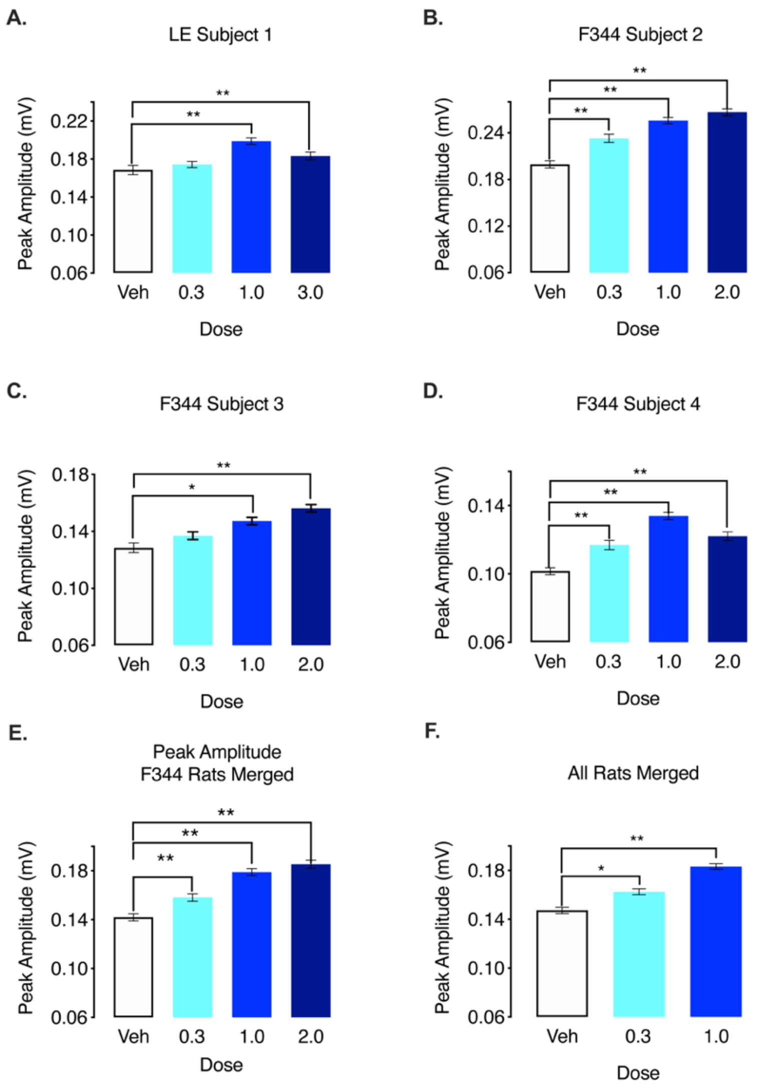


**Fig. S16.** Oral treatment with $\alpha$5IA dose-dependently increases peak amplitude of ripples in adult male rats. **A)** Results for LE Subject 1 showing significant dose-dependent increases in average peak amplitude of ripple events following administration of 1.0 and 2.0 mg/kg doses of $\alpha$5IA. **B)** Histogram for F344 Subject 2 showing significant dose-dependent increases in average peak amplitude of ripple events at all three doses of $\alpha$5IA tested. **C)** Histogram for F344 Subject 3 showing significant dose-dependent increases in average peak amplitude of ripple events following administration of 1.0 and 2.0 mg/kg doses of $\alpha$5IA. **D)** Histogram for F344 Subject 4 showing significant dose-dependent increases in average peak amplitude of ripple events following administration of $\alpha$5IA. **E)** Average peak amplitude of ripples from all three F344 rats demonstrates significant dose dependent increases of 15% (0.3 mg/kg), 31% (1.0 mg/kg), and 20% (2.0 mg/kg). **F)** Histogram showing average peak amplitude of ripples from all four rats tested demonstrates significant dose dependent increases of 15% (0.3 mg/kg), 31% (1.0 mg/kg), and 20% (2.0 mg/kg). Significance indicated by * at p < 0.01 and ** at p < 0.001.

##### *3. Effects of $\alpha$5IA on Ripple Duration in Wild Type Rats*

Kruskal-Wallis ANOVA revealed no consistent significant within subject effects of drug on ripple duration. Subject 1 (LE Rat; age 15 mo: #327396; showed no significant drug induced effects on ripple duration ($\chi^2$ (3, 104) = 1.0; p = 0.78). (**Fig. S17-A**) For Subject 2 (F344 Rat; age 9 mo: #130) the ANOVA was significant ($\chi^2$ (3, 104) = 8.5 p = 0.036), but the pairwise comparisons of each dose with vehicle did not reveal any significant changes in ripple duration. (**Fig. S17-B**) A significant effect of drug on ripple duration is observed following administration of 1.0 mg/kg α5IA in Subject 3 (F344 Rat age 16 mo: #600040; ($\chi^2$ (3, 104) = 3.9; p = 0.002). (**Fig. S17-C**) A significant effect of drug on ripple duration is observed in Subject 4 (F344 Rat age 12 mo: #113; $\chi^2$ (3, 160) = 4.4; p = 0.04) on a planned comparison at the 0.3 mg/kg dose but this was not after controlling for multiple comparisons ($\chi^2$ (3, 160) = 4.4; p = 0.07). (**Fig. S17-D**). Despite the lack of any consistent within subject changes in ripple duration, ANOVA of the merged ripple duration data from all three F344 rats revealed a small but significant ($\chi^2$ (3, 405) = 10.4; p = 0.04) 11% decrease following administration of the 1.0 mg/kg dose of α5IA. (**Fig. S17-E**) A significant ($\chi^2$ (3, 509) = 10.4; p = 0.018) decrease in ripple duration of 10% is also seen when the data from all for four animals including Subject 1 (LE Rat age 15 mo: # 327396) are merged prior to analysis (**Fig. S17-F**).

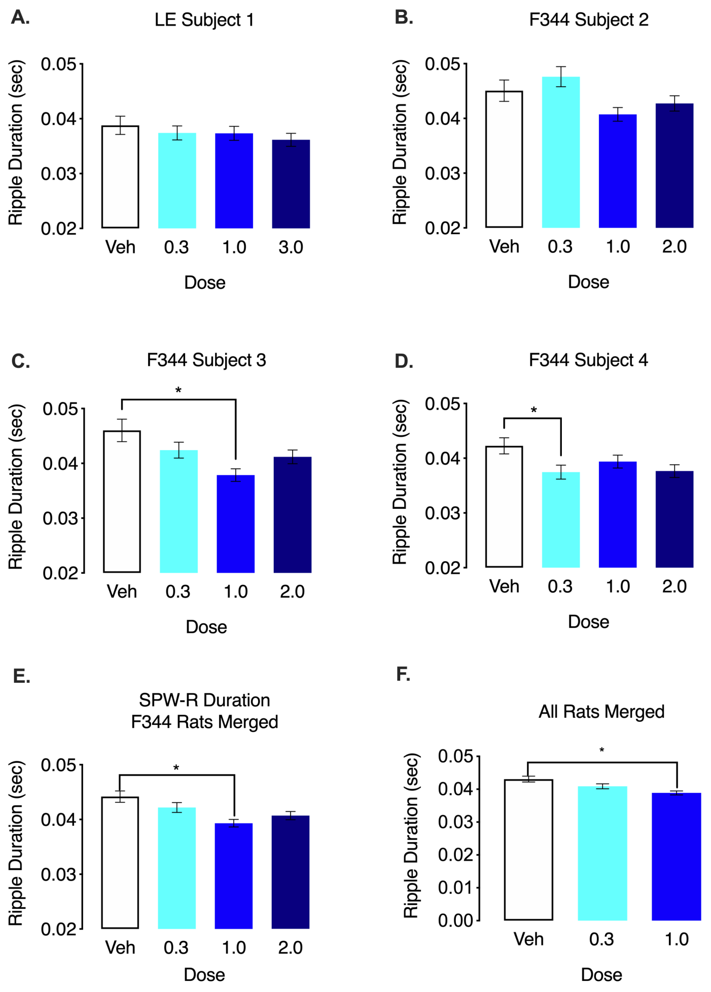


**Fig. S17.** Oral treatment of adult wild type rats with 1.0 mg/kg α5IA decreases the average duration of ripple events. **A)** Subject 1 (LE Rat #600040) shows no significant dose-dependent changes in average ripple duration following administration of α5IA. **B)** Subject 2 (F344 Rat #130) shows no significant changes in average ripple duration. **C)** Subject 3 (F344 Rat #600040) shows a significant decrease in ripple duration following 1.0 mg/kg α5IA. **D)** Subject 4 (F344 Rat #113) shows a significant decrease in average ripple duration following administration of 0.3mg/kg α5IA. **E)** Merged data from all three F344 rats reveals a significant decrease in ripple duration following administration of 1.0 mg/kg dose of α5IA. **F)** A significant 10% decrease in ripple duration is seen following administration of 1.0 mg/kg dose of α5IA in the merged data from all 4 wild type rats. Significance indicated by * at $p < 0.05$.

*4. Effects of α5IA on Peak Frequency of Ripples in Wild Type Rats*

No significant dose-dependent changes in peak ripple frequency are observed following treatment of wild type rats with α5IA (**Data not shown**).

5. *Supplemental Results for Effects of α5IA on Ripple Band Power in TgF344-AD Rats*

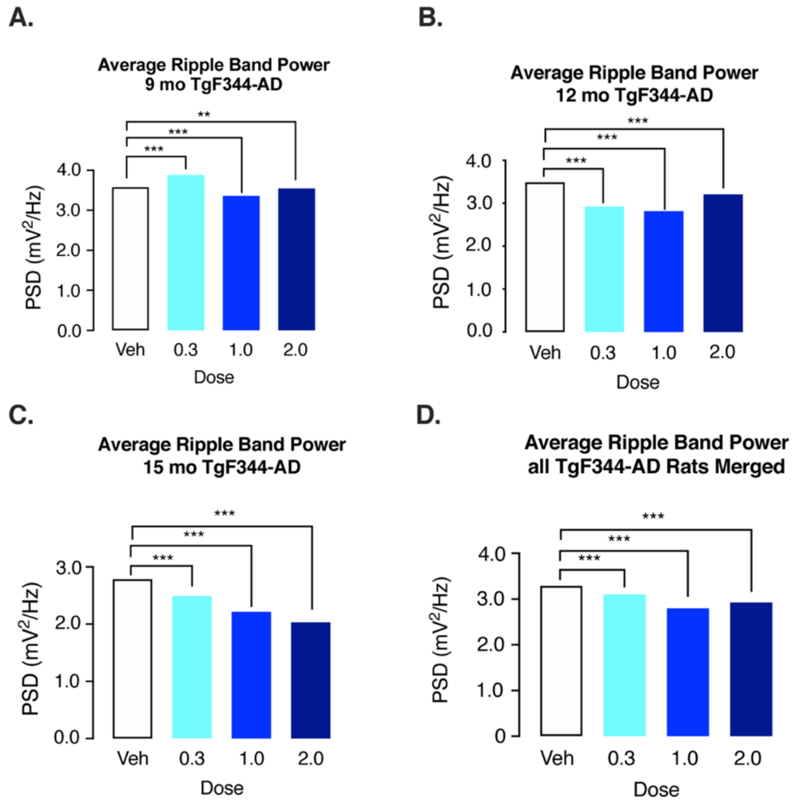


**Fig S18**: Treatment of adult TgF344-AD rats with α5IA decrease average power in ripple band. **A)** Data from 9 mo TgF344-AD shows increase at 0.3 mg but a decrease in average ripple band power following 1.0 and 2.0 mg/kg α5IA. **B)** Analysis of ripple band power in 12 mo TgF344-AD shows significant decrease in ripple band power following all 3 doses of α5IA. **C)** Analysis of ripple band power in 15 mo TgF344-AD shows significant decrease in ripple band power following all 3 doses of α5IA. **D)** Merged data from all three F344 rats reveals a significant decrease in ripple band power following administration of all three doses of dose of α5IA. Significance indicated by *** at $p < 0.001$.
